# TCR signal strength during positive selection shapes CD25 expression patterns on thymically derived regulatory T cells

**DOI:** 10.64898/2026.06.16.732736

**Authors:** Isabel Baldwin, Dunfang Zhang, Yash Kulkarni, Wanjun Chen, Ellen Robey

**Affiliations:** University of California, Berkeley, Division of Immunology and Molecular Medicine, Department of Molecular and Cell Biology, Berkeley, CA; Mucosal Immunology Section, National Institute of Dental and Craniofacial Research, National Institutes of Health, Bethesda, MD, United States

## Abstract

Regulatory T cells (Tregs) are a suppressive subset of CD4 T cells that maintain immune homeostasis. Most Tregs develop in the thymus but how thymic selection impacts peripheral Treg fate remains unknown. Here we show that the strength of TCR signal experienced during positive selection in the thymic cortex impacts how CD4 SP thymocytes respond to Treg inducing signals in the medulla. Thymocytes that experienced weak signals during positive selection (identified as CD5^LO^ CD4 SP) show a greater dependence on TGFβ than IL-2 for Treg induction and produce Foxp3+ cells that lack constitutive expression of the classic Treg marker CD25. Moreover, CD4 clones that give rise to CD25- thymic Tregs also produce CD25- peripheral Tregs. These data indicate that positive selection provides critical context for how CD4 T cells respond to TCR agonist and cytokine signals, leading to an alternative lineage of Tregs that lacks constitutive CD25 expression.

## Introduction

Foxp3-expressing regulatory T cells (Tregs) are an immunosuppressive subset of CD4+ T cells that are required for maintaining tolerance to self (Sakaguchi *et al*., 2020). Tregs exhibit considerable heterogeneity, differing in their state of activation, memory vs. effector-like properties, and location in tissues vs. secondary lymphoid organs (SLO) (Campbell and Koch, 2011). Most of the stable peripheral Treg pool is thymically derived (Sawant and Vignali, 2014), and is generated when CD4+CD8- thymocytes (single positive or CD4 SP) encounter agonist self-ligand in the thymus. The impact of thymic selection on peripheral Treg heterogeneity remains largely unknown.

During development, thymocytes undergo two spatially and temporally distinct waves of screening for T cell receptor (TCR) self-reactivity. Thymocytes first undergo positive selection in the thymic cortex, where intermittent contacts with self-peptide/MHC complexes provide survival signals and “tune” the thymocyte’s responsiveness to self peptide-MHC complexes displayed in the cortex. The cell surface protein CD5 is part of the tuning apparatus and serves a robust marker for the self-reactivity of mature thymocytes and T cells (Azzam *et al*., 1998, 2001). Positive selection tuning has recently been appreciated as an important factor shaping the responsiveness of naïve conventional T cells to antigen and cytokines. (Cho *et al*., 2010; Palmer *et al*., 2011; Mandl *et al*., 2013, 2026; Persaud *et al*., 2014; Fulton *et al*., 2015; Eggert and Au-Yeung, 2021). After positive selection thymocytes migrate from the thymic cortex to the medulla, where they are faced with a diverse new set of tissue-restricted self-peptides, partly due to AIRE-mediated promiscuous gene expression by medullary thymic epithelial cells (Miller *et al*., 2024). Strong recognition of medullary self-peptides by a CD4 SP thymocyte can lead to either death by negative selection, upregulation of the transcription factor Foxp3 to adopt a Treg fate or escape of autoreactive CD4 T cells to the periphery. While there is evidence that positive selection tuning impacts Treg induction in the periphery (Henderson *et al*., 2015; Pennock *et al*., 2025), its impact on thymic Treg development remains unknown.

In addition to TCR triggering by agonist self-peptide, cytokine signals via the IL-2/STAT5 pathway are also critical for thymic Treg development (Fontenot *et al*., 2005; Soper, Kasprowicz and Ziegler, 2007; Vang *et al*., 2008). There is evidence that a strong TCR signal causes CD4 SP thymocytes to upregulate CD25 (the high affinity IL-2 receptor chain), generating a CD25+Foxp3- precursor population. Treg precursors that subsequently receive IL-2 -induced STAT5 signaling can turn on Foxp3 expression (Burchill *et al*., 2007, 2008; Lio and Hsieh, 2008). There is evidence that transforming growth factor beta (TGFβ), which has a well-established role in Treg induction in the periphery (Chen *et al*., 2003; Kretschmer *et al*., 2005), also plays a role in thymic Treg development (Liu *et al*., 2008; Chen and Konkel, 2015). How IL-2 and TGFβ work together to promote thymic Treg development, and whether all thymic CD4 SP have an equal dependence on these two cytokine pathways to drive Treg development remains unclear.

Prior to the discovery of Foxp3 as the master transcription factor defining Treg identity, CD25 was the primary marker used to identify Tregs (Takahashi *et al*., 1998). However, it is an imperfect marker, since conventional T cells transiently express CD25 upon activation, and not all Foxp3+ CD4 T cells express CD25 (Apostolou *et al*., 2002; Zelenay *et al*., 2005; Ono *et al*., 2006; Coleman *et al*., 2012). Moreover, a subset of thymic Foxp3+ cells lack CD25 (Marshall *et al*., 2014; Owen *et al*., 2019; Apert *et al*., 2022), in apparent contradiction of the notion that IL-2/STAT5 signaling is needed to induce Foxp3. It has been proposed that CD25- thymic Tregs are alternative precursors to CD25+ thymic Tregs (Marshall *et al*., 2014; Owen *et al*., 2019). However, the extent to which CD25-Foxp3+ thymocytes maintain their CD25- phenotype after they leave the thymus has not yet been addressed.

Here we use TCR transgenic models, a thymic slice system of Treg development, and TCR repertoire analyses to address these questions. We show that positive selection tuning impacts CD25 expression by thymically derived Tregs, such that CD4 SP with low cortical self-reactivity (CD5^LO^) largely give rise to CD25- thymic Tregs while CD4 SP with moderate cortical self-reactivity (CD5^HI^) tend to produce CD25+ thymic Tregs. We provide evidence that positive selection impacts the relative dependence on IL-2 versus TGFβ for Treg induction; CD5^LO^ CD4 SPs are more TGFβ dependent while CD5^HI^ CD4 SPs are more IL-2 dependent. We also provide genetic evidence that CD25- thymic Tregs are more dependent on TGFβ for their development *in vivo*. Finally, we show that CD4 clones that develop into CD25- thymic Tregs remain CD25- after they leave the thymus. Our data suggest that positive selection tuning in the thymic cortex provides critical context for how CD4 T cells respond to TCR agonist and cytokine signals in the medulla, such that CD4 SP that experience relatively weak signals during positive selection followed by encounter with agonist ligands in the medulla produce an alternative lineage of Tregs that lacks constitutive CD25 expression.

## Results

### TCR transgenic thymocytes and thymic slice culture recapitulate CD25+ and CD25- pathways for Treg development

To investigate how TCR specificity influences thymic Treg CD25 expression, we used a thymic slice culture system in which CD4 SP thymocytes encounter agonist ligands and undergo a wave of Treg development *in situ* (Weist *et al*., 2015). Briefly, we overlaid thymocytes from mice expressing rearranged MHC-II specific TCR transgenes onto thymic slices along with bone marrow derived dendritic cells (BMDCs) presenting the corresponding agonist peptide **(Fig. 1A)**. We used congenically distinct or CellTrace-labeled thymocytes for overlaid populations, allowing us to separate them from slice resident thymocytes by flow cytometry **(Suppl. Fig. 1).** We used three TCR transgenic models: OTII (Barnden *et al*., 1998), which recognizes a peptide from chicken ovalbumin (OVAp); BDC2.5 (Katz *et al*., 1993), which recognizes a hybrid insulin peptide (2.5HIP); and AND (Hedrick *et al*., 1982), which recognizes a peptide from moth cytochrome c (MCCp) (**Table 1**). After 72 hours (72h) of culture, we assessed Treg development by the appearance of TCR transgenic Foxp3+ CD4 single positive (CD4 SP) thymocytes. While agonist peptide led to the development of Foxp3+ Tregs for all three TCR transgenic models **(Fig. 1B,C)**, the proportions of CD25+ and CD25- Tregs varied **(Fig. 1D).** Specifically, OTII produced predominantly CD25+Tregs, BDC2.5 produced a large proportion of CD25-Tregs, and AND produced a mix of CD25+ and CD25- Tregs **(Fig.1D**). Thus, three different TCR transgenic lines exhibit distinct propensities to produce CD25- or CD25+ Tregs.

**Figure 1.**
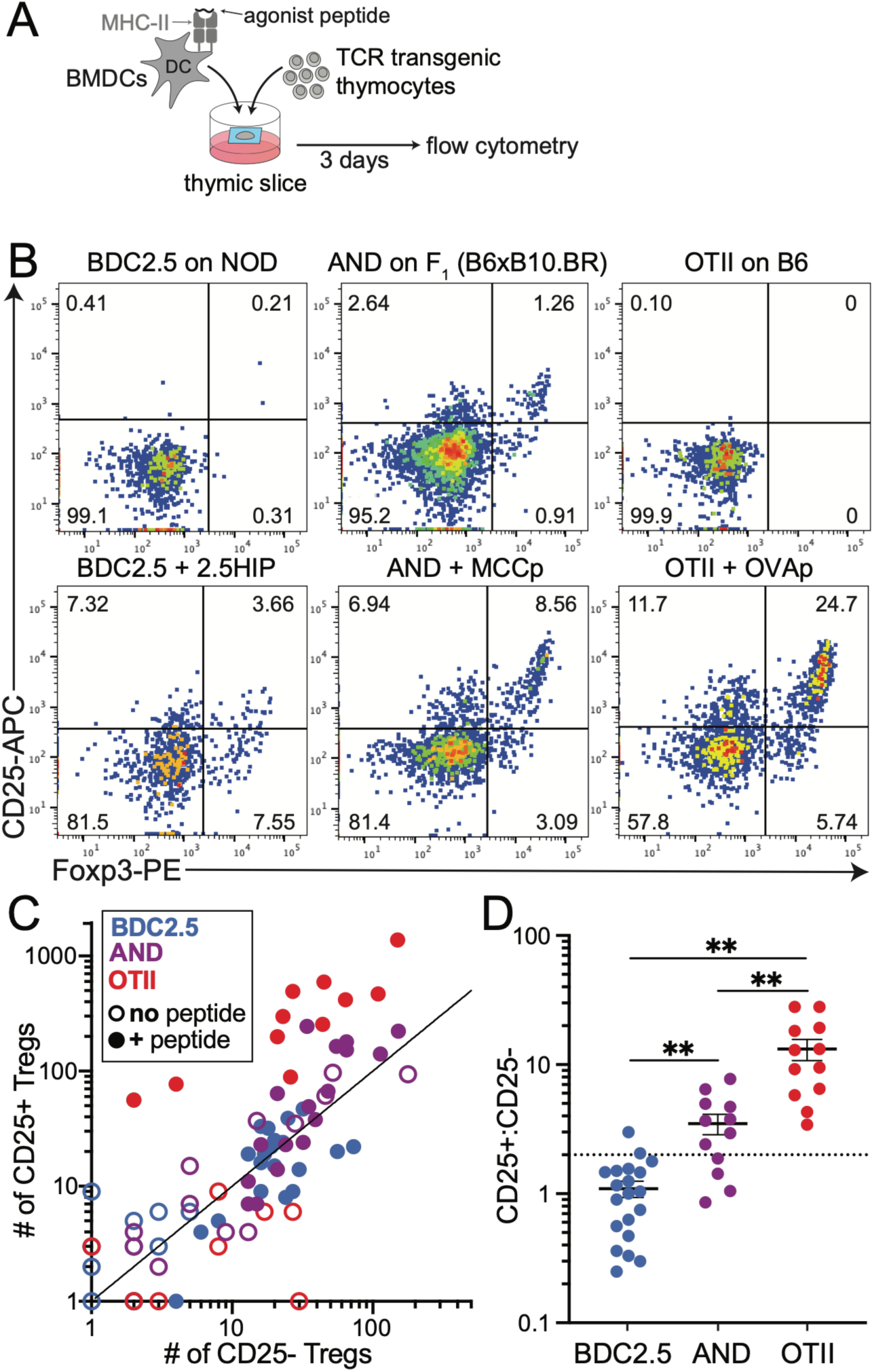
TCR transgenic thymocytes and thymic slice culture model different pathways of thymic Treg development. A. Experimental schematic. Thymocytes from 3 different TCR transgenic models (BDC2.5, AND, or OTII) were cultured on thymic tissue slices along with bone marrow dendritic cells (BMDCs) loaded with their corresponding agonist peptides (Table 1) and analyzed by flow cytometry after 72 hours. Thymic slices and BMDC were from NOD mice for the BDC2.5 thymocytes, from F_1_ (B6xB10.BR) mice for AND, and from B6 mice for OTII. B. Representative flow plots showing CD25 and Foxp3 expression on gated CD4+CD8- thymocytes of TCR transgenic donor origin from slice cultures. C-D. Compiled data. Each symbol represents one thymic slice from 3 separate experiments with 3-4 slices per condition. C. Total number of CD25+ and CD25- Foxp3+ Tregs recovered from each thymic slice under the indicated conditions. D. Ratio of CD25+ Foxp3+ over CD25- Foxp3+ Tregs for samples with agonist peptide. Horizontal dotted line represents the average ratio of CD25+: CD25- Tregs from WT slice resident CD4 SP. Horizontal lines show average +/- SEM, and **=P<0.01 (one way analysis of variance (ANOVA) with multiple comparisons).

**Table 1.**
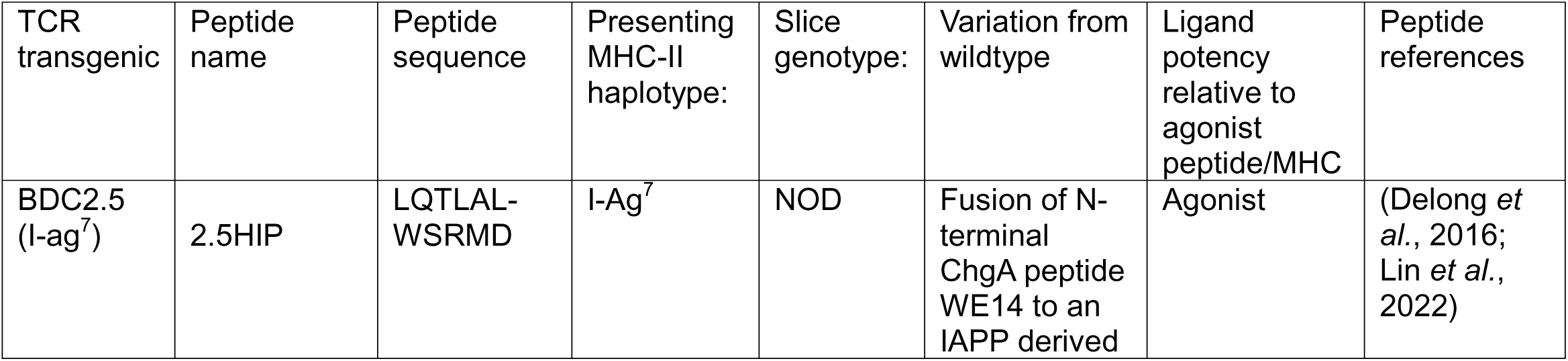
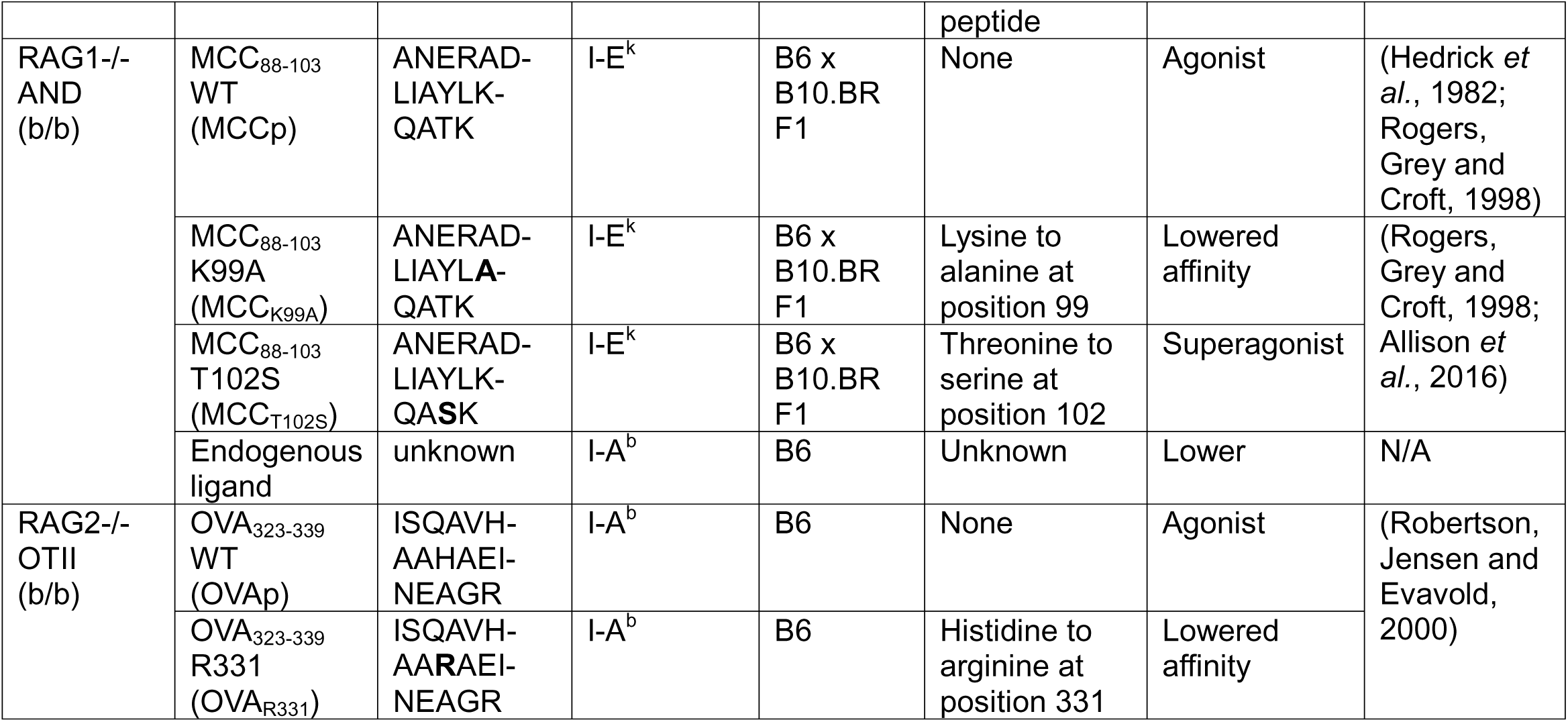
Summary of agonist peptides used in thymic Treg development slice experiments.

### TCR signal strength during both positive selection and agonist selection influence the pathway for Treg development

Previous studies provided evidence that CD25-thymic Treg experience weaker TCR signals compare to CD25+ Tregs (Owen *et al*., 2019), but whether this reflects weaker signals during positive selection in the cortex, during tolerance induction in the medulla, or both remains unclear. Indeed, given evidence that TCR signal strength during positive selection impacts T cell response to agonist ligands (Mandl *et al*., 2013; Persaud *et al*., 2014; Fulton *et al*., 2015), it seemed plausible that functional tuning during positive selection might underlie the different responses of the three TCR transgenic models to agonist TCR ligands in thymic slice culture. To investigate this question, we assessed the expression of CD5, which provides a surrogate marker of self-reactivity on mature thymocytes and naïve T cells (Azzam *et al*., 1998, 2001). Interestingly, CD5 levels on CD4 SP from BDC2.5 TCR transgenic mice (BDC2.5 CD4 SP) were at the extreme low end of the distribution observed for polyclonal CD4 SP from B6 wildtype (WT) mice, while CD4 SP from OTII TCR transgenic mice (OTII CD4 SP) were at the higher end **(Fig. 2A).** Thus, the ability of three TCR transgenic models to produce CD25+ Tregs upon agonist selection positively correlated with their cortical self-reactivity.

**Figure 2.**
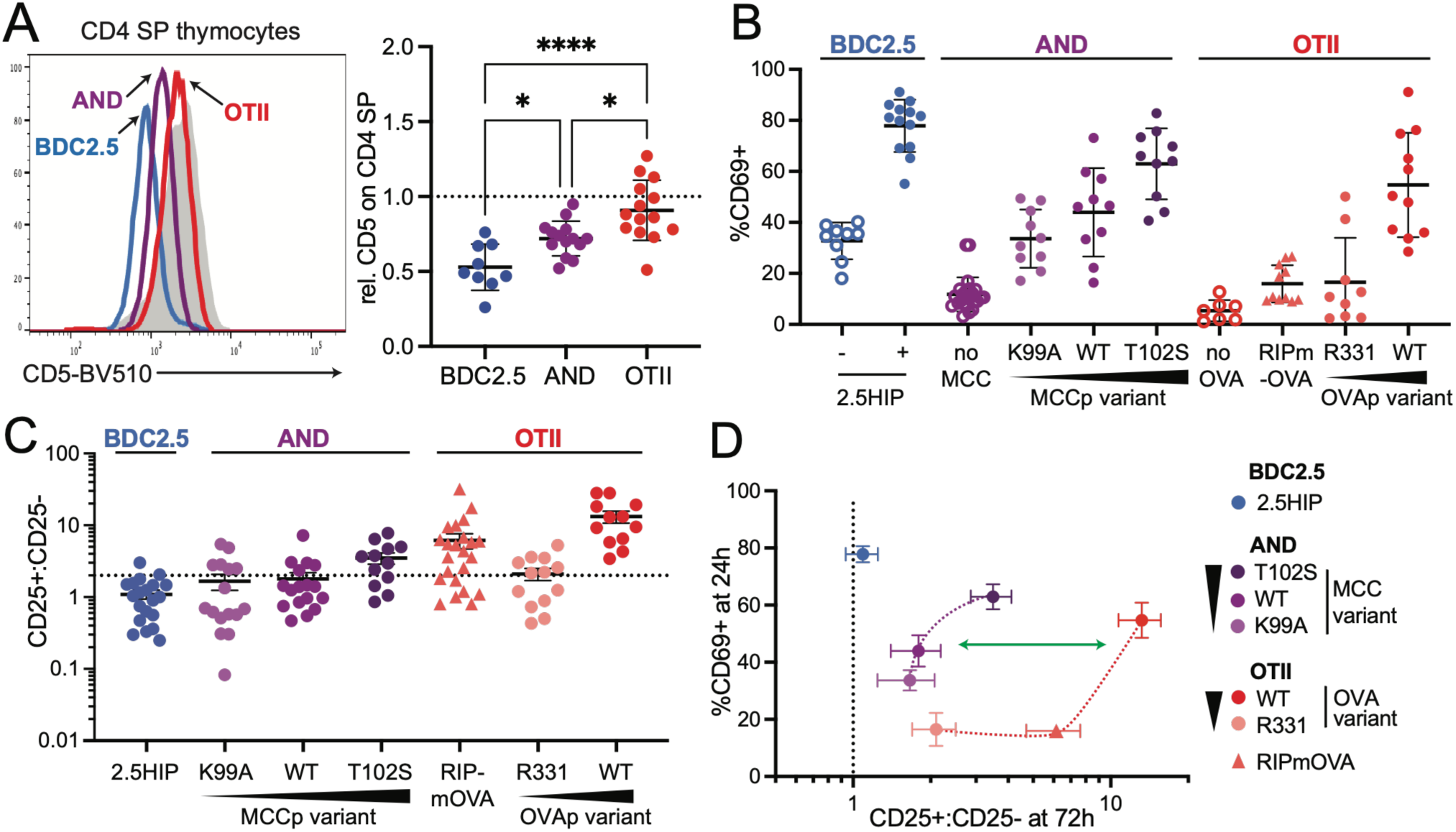
TCR signal strength during both positive and agonist selection influence the pathway for Treg development. A. CD5 levels on mature CD4 SP thymocytes (gated CD8-CD4+CD69-) from the indicated TCR transgenic mice were determined by flow cytometry. Gray histograms are CD5 levels on mature CD4 SP thymocytes from B6 WT. Plots show representative flow cytometry (left) or compiled data (right). CD5 gMFI for each sample was normalized to CD5 levels on equivalent WT cell population analyzed in the same experiment. Six separate experiments with 1-2 mice each were performed, and each dot represents an individual mouse. Average values are indicated by horizontal bars and range marks indicate SD. For statistical analysis, * = P <0.05, ****=P<0.0001 (one way analysis of variance (ANOVA) with multiple comparisons). B-D. BDC2.5, AND, or OTII thymocytes were cultured on thymic slices along with peptide-loaded BMDC for 24 hours to assess TCR activation level by CD69 induction (A,C) or for 72 hours to assess CD25+ and CD25- Treg development (B, C). Open circles indicate BMDC with no peptide, and closed circles indicate BMDC loaded with the indicated peptide (Table 1). OTII thymocytes overlaid onto thymic slices from transgenic mice with AIRE-dependent expression of OVA (Kurts 1996) (triangles) were included for comparison. For AND thymocytes with MCC peptide variants, BMDC and thymic slices were from F_1_ (B6xB10.BR) mice. Data are from at least 3 separate experiments with 3-4 slices per condition. Small horizontal and vertical bars represent the SEM. A, B show compiled data for the indicated TCR transgenic model and antigen condition with each dot representing one thymic slice. C shows data from A and B replotted to emphasize the relationship between TCR activation (% CD69+ at 24 hours (Y axis) with Treg development (CD25+ to CD25- Treg ratio at 72 hours (X axis). Each dot represents the mean of all replicates for the indicated experimental condition.

To separately assess the role of agonist peptide affinity on CD25 expression by thymic Tregs, we cultured TCR transgenic thymocytes on thymic slices together with BMDCs loaded with peptides of varying agonist potencies **(Table 1).** We read out expression of the TCR target gene CD69 after 24h of culture, a time point of maximal induction by agonist peptides across different conditions **(Suppl. Fig. 2A)**. As expected, CD69 induction after 24h correlated with known ligand potency for the OTII and AND TCR transgenic models **(Fig. 2B, black triangles, Suppl. Fig. 2A)**. Quantification of CD25+ and CD25- Foxp3+ Treg development after 72h of culture revealed that AND Tregs exhibited greater CD25 expression when induced by a higher affinity ligand and lower expression when induced by a lower affinity peptide. In addition, OVA_R331_ (a variant OVA peptide with lower affinity, **Table 1**) produced OTII Tregs with reduced CD25 expression **(Fig. 2C, black triangles).** Thus, for an individual TCR, higher agonist signal strength tends to correlate with CD25 expression by thymic Tregs.

To compare the relationship between agonist signal strength and Treg phenotype across the three TCR transgenic models, we plotted CD69 expression in transgenic CD4 SP thymocytes at 24h (as a measure of the strength of TCR response to agonist ligands) versus the ratio of CD25+ to CD25- Treg present at 72h (**Fig. 2D**). This plot reveals both the impact of agonist ligand potency within each TCR transgenic model, as well as the inherent tendency of each TCR transgenic model to produce CD25+ or – Tregs. For example, AND thymocytes exposed to the superagonist peptide MCC_T102S_ have equivalent CD69 induction as OTII thymocytes exposed to OVAp but have a significantly lower ratio of CD25+ to CD25- Tregs under these conditions (**Fig. 2D**, green arrow). In addition, exposure of BDC2.5 thymocytes to 2.5HIP peptide led to the highest CD69 expression at 24h but one of the lowest ratios of CD25+ to CD25- Tregs at 72h. Taken together, these data suggest that both positive selection in the cortex and agonist selection in the medulla influence CD25 expression by thymic Tregs.

### The AND TCR recognizes an endogenous agonist TCR ligand that can drive thymic Treg development

Thymocytes naturally encounter distinct sets of self-peptides during positive vs. negative selection due to the unique peptide repertoire of cortical thymic epithelial cells and the presence of diverse tissue restricted antigen presentation in the medulla (Klein *et al*., 2014; Ohigashi, Matsuda-Lennikov and Takahama, 2021; Mandl *et al*., 2026). As a result, an individual TCR may confer low self-reactivity during positive selection while also exhibiting strong reactivity to medullary and peripheral self-peptides. The presence of a small but consistent population of Foxp3+ AND CD4 SP in thymic slice culture in the absence of MCC peptide **(Fig. 1B, C)**, suggests that the AND TCR may be an example of a TCR that reacts with an endogenous high affinity self-ligand found in the thymic medulla, despite its low cortical self-reactivity. Although AND TCR rag-/- mice have no detectable thymic Tregs, we considered that clonal competition may impair their development, as reported previously (Bautista *et al*., 2009; Leung, Shen and Lafaille, 2009). To test this, we compared the Treg development of AND thymocytes overlaid onto B6 WT slices (where AND frequency was <5%) to AND thymocytes overlaid onto thymic slices prepared from AND TCR transgenic mice (where AND frequency was 100%). We observed Treg development at 72h on B6 WT, but not on AND slices **(Suppl. Fig. 3A).** We also observed that AND thymocytes cocultured *in vitro* with BMDCs without added peptide showed a small, but significant increase in CD69 above the levels observed without BMDCs, while this was not the case for OTII thymocytes **(Suppl. Fig. 3B, C**). All together these data indicate that the AND TCR recognizes an unknown agonist ligand in B6 WT mice that can drive Treg development when the frequency of AND thymocytes is low. In this study we use both exogenous MCCp-loaded BMDCs and the unknown endogenous ligand as alternative ways of experimentally generating AND Tregs.

While CD5 levels are primarily set during positive selection (Persaud *et al*., 2014), chronic exposure to tolerizing peripheral antigens can lead to a gradual upregulation of CD5 on mature T cells (Stamou *et al*., 2003; Hawiger *et al*., 2004). Interestingly, while CD5 levels on thymic AND CD4 SP were lower than WT (**Fig. 2A**), splenic AND T cells exhibited higher CD5 expression compared to WT CD4 T cells (**Suppl. Fig. 3D**) (Zinzow-Kramer, Weiss and Au-Yeung, 2019). This implies that low CD5 expression on AND thymocytes established during positive selection remains stable until after AND T cells leave the thymus and repeatedly encounter agonist ligand in the periphery. Additionally, we noted that CD5 expression levels did not change significantly after 72h of exposure to agonist ligands in thymic slice culture, and that the relative CD5 expression levels on CD4 SP and Tregs between TCR transgenic models (BDC2.5<AND<OTII) was retained **(Suppl. Fig 3E)**. Altogether, these data indicate that, although prolonged encounter with agonist ligand can lead to a gradual increase in CD5 expression on peripheral T cells, the CD5 levels on thymic CD4 SP provide a robust readout of positive selection signal strength, even for thymocytes that encounter high affinity agonist ligands in the medulla.

### Niche-localized IL-2 production does not alter CD25 expression on AND Tregs

CD5^HI^ vs. CD5^LO^ CD4 SP thymocytes have been reported to differ in their ability to produce IL-2 upon TCR stimulation (Persaud *et al*., 2014; Zinzow-Kramer, Weiss and Au-Yeung, 2019). To confirm this observation for the three TCR transgenic models in the current study, we stimulated BDC2.5, AND, and OTII thymocytes with PMA and ionomycin and quantified IL-2 production in gated CD4 SP thymocytes after 16h. As expected, IL-2 production positively correlated with CD5 expression on TCR transgenic CD4 SP, with OTII CD4 SP producing the most IL-2 and BDC2.5 the least **(Fig. 2A**, **Fig. 3A)**, We saw the same IL-2 production trend (OTII>AND>BDC2.5) when we stimulated TCR transgenic thymocytes with agonist peptide-loaded BMDCs, in spite of similar TCR stimulation, as read out by induction of CD69 **(Suppl. Fig. 4A).** In addition, we could detect IL-2 production by OTII CD4 SPs after 24h of culture in thymic slice culture with OVAp, but not by AND or BDC2.5 CD4 SPs in slices with their high affinity peptides **(Fig. 3B).**

**Figure 3.**
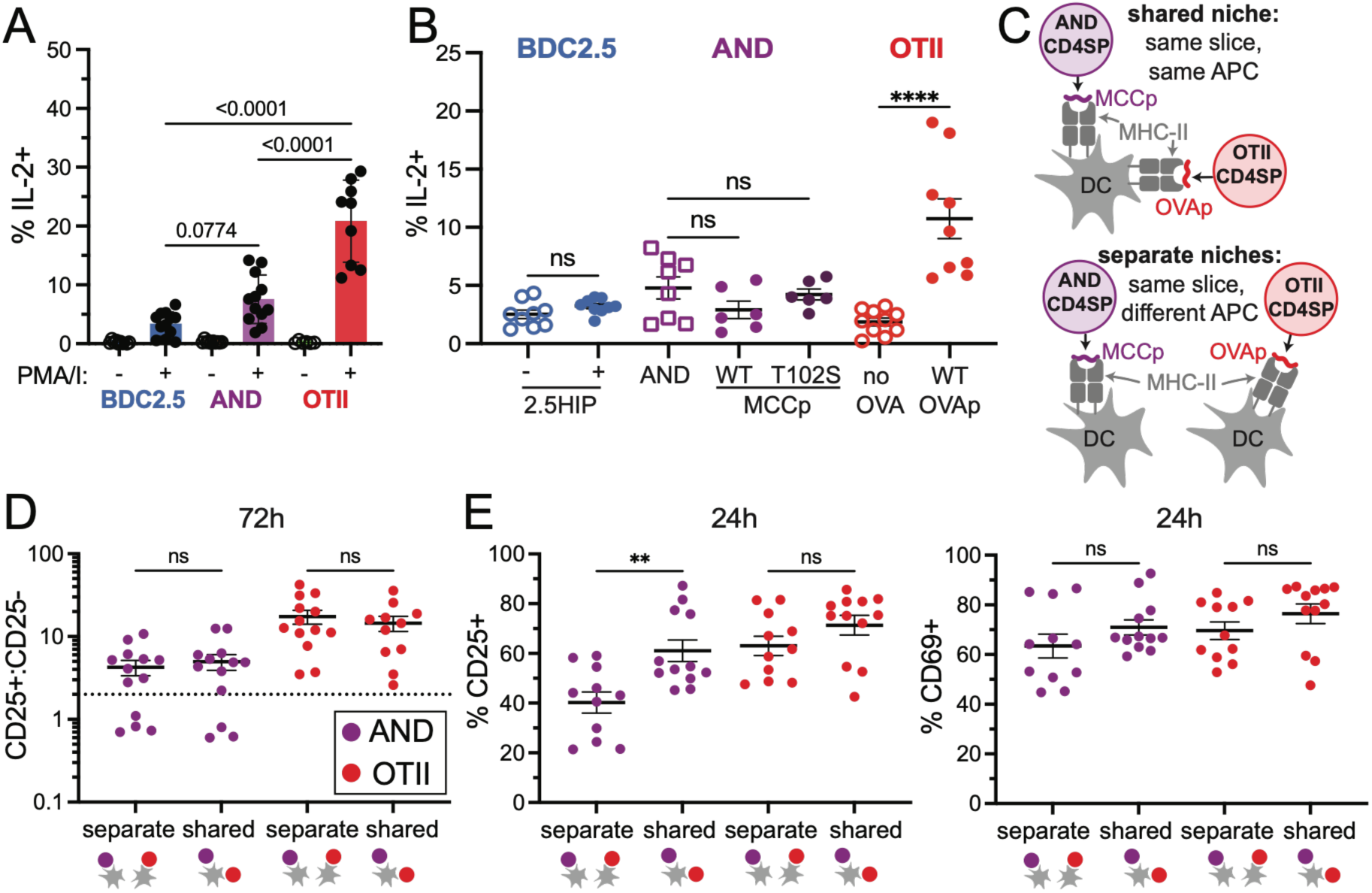
Niche-localized IL-2 production does not alter CD25 expression on AND Tregs. A,B. TCR transgenic thymocytes were stimulated *in vitro* with PMA and ionomycin for 16 hours (A) or cultured in thymic slices with BMDC loaded with the indicted agonist peptide for 24 hours (B), and IL-2 expression by TCR transgenic CD4 SP was determined by flow cytometry. Data is pooled from at least three experiments, and each symbol represents a replicate sample. C. Experimental design. In the shared niche condition (top), BMDC present both OVAp and MCCp, promoting interactions of AND and OTII thymocytes with the same APC. In the separate niche condition (bottom), BMDCs present either OVAp or MCCp, so AND and OTII thymocytes interact with different APC within the thymic slice. D, E. AND and OTII thymocytes were cultured on thymic slices along with peptide loaded BMDC from F1 (B6 x B10.BR) mice. For shared niche conditions BMDC were incubated with both OVA and MCC peptides. For separate niche conditions BMDC were separately loaded with either OVAp or MCCp and then mixed before adding to thymic slices. Both AND and OTII thymocytes were added to each thymic slice, and samples were analyzed after 72 hours to assess CD25+ and CD25- Treg development (D) or 24 hours to assess CD69 and CD25 induction (E) from AND or OTII donor populations. Each symbol represents one thymic slice from at least 3 separate experiments with 3-4 replicates per condition. Data is represented as the average +/- SEM. For statistical analysis, one-way ANOVA with multiple comparisons was performed. **=P<0.01, ****=P<0.0001

Given evidence that Treg development depends on IL-2 produced by self-reactive CD4 SPs (Owen *et al*., 2018; Hemmers *et al*., 2019), and that IL-2 is required in close proximity to the agonist peptide presenting cell (Weist *et al*., 2015), we considered that IL-2 produced by agonist signaled OTII thymocytes in the thymic slice culture system may impact the development of nearby Treg precursors. Specifically, since IL-2 mediated STAT5 signaling can increase CD25 expression (Smith and Cantrell, 1985; Kim and Leonard, 2002), we considered that IL-2 production by OTII CD4 SP could drive higher CD25 expression on Treg precursors that interact with the same APC.

To ask how APC-associated IL-2 affects CD25 expression on thymic Tregs, we designed an experiment to test whether IL-2 produced by OTII CD4 SP thymocytes could increase CD25 expression on AND Tregs when they interact with the same APC (**Fig. 3C)**. We incubated BMDCs with <u>both</u> MCCp and OVAp and overlaid them onto thymic slices along with AND and OTII thymocytes (“shared niche” condition). For comparison, we incubated BMDCs separately with <u>either</u> MCCp or OVAp and overlaid them, along with AND and OTII thymocytes, onto the same thymic slices (“separate niche” conditions, **Fig. 3C**). As an additional control, we also cultured AND and OTII thymocytes separately on slices along with BMDCs loaded with their corresponding agonist peptides (“separate slice” conditions) **(Suppl. Fig. 4B).** We then assessed AND and OTII Foxp3+ Treg development under different conditions after 72h. OTII and AND Tregs exhibited a similar ratio of CD25+ to CD25- Tregs regardless of whether they developed in a shared or separate niche (**Fig. 3D, Suppl. Fig. 4C,D).** On the other hand, at 24h AND CD4 SP in a shared niche with OTII CD4 SP expressed higher CD25 than AND CD4 SP in a separate niche **(Fig. 3E)**. The increase in CD25 for AND CD4 SP in the shared niche was likely due to increased IL-2/STAT5 rather than TCR signaling, because these conditions did not lead to a significant increase in CD69 expression **(Fig. 3E).** AND and OTII thymocytes showed similar CD25 and CD69 induction in separate niches conditions compared to when they were cultured on separate slices (**Suppl. Fig. 4B,E,F)**, confirming the lack of interaction between the DC-associated niches under the separate niche conditions. These data suggest that increased IL-2 production from OTII CD4 SPs can drive transient CD25 upregulation on neighboring AND CD4 SPs but is not sufficient to override their propensity to develop as CD25- Tregs.

### CD4 SP with moderate vs low cortical self-reactivity have distinct requirements for Treg-inducing cytokines

Given evidence that niche-associated IL-2 levels did not determine CD25 expression by thymic Tregs, we considered whether thymocyte intrinsic differences imposed by functional tuning during positive selection might impact how CD4 SP cells respond to Treg inducing cytokines (Cho *et al*., 2010; Palmer *et al*., 2011; Fulton *et al*., 2015). To test this, we examined the ability of CD5^HI^ vs. CD5^LO^ mature CD4 SP to convert into Foxp3-expressing Tregs *in vitro* upon stimulation with CD3/CD28 antibodies in the presence or absence of IL-2 and/or TGFβ (**Fig. 4A-C**). For these experiments, we used non-TCR transgenic Foxp3^RFP^ reporter mice, which exhibit a broader distribution of CD5 compared to TCR transgenic CD4SP, and allows us to sort for Foxp3- (RFP negative) CD4 SP as well as the top and bottom 25% of CD5 expression prior to *in vitro* stimulation (**Fig. 4A**). While both CD5^HI^ and CD5^LO^ populations gave rise to Foxp3^RFP^+ Tregs with similar high efficiency when <u>both</u> IL-2 and TGFβ were added to the cultures, distinct responses were observed in the presence of either IL-2 or TGFβ (**Fig. 4B**). Specifically, IL-2 was most effective at inducing Foxp3 expression in CD5^HI^ CD4 SP, while TGFβ was the more potent Foxp3 inducer for CD5^LO^ CD4 SP. To explore the impact of endogenous IL-2 and TGFβ produced by CD4 SP during culture, we also performed *in vitro* cultures in presence of IL-2 or TGFβ blocking antibodies. We observed that blocking TGFβ in the presence of added IL-2 greatly reduced Treg development from both populations **(Fig. 4B**). Likewise, when we blocked IL-2 in the presence of added TGFβ, Treg development from both CD5^HI^ and CD5^LO^ CD4 SP was barely detectable **(Fig. 4B**). Taken together, these results indicate that, while both cytokines are needed for optimal Treg development, IL-2 is a more important inducer of Tregs for CD4 SP with moderate cortical self-reactivity (CD5^HI^), whereas TGFβ is a more important Treg inducer for CD4 SP with low cortical self-reactivity (CD5^LO^).

**Figure 4.**
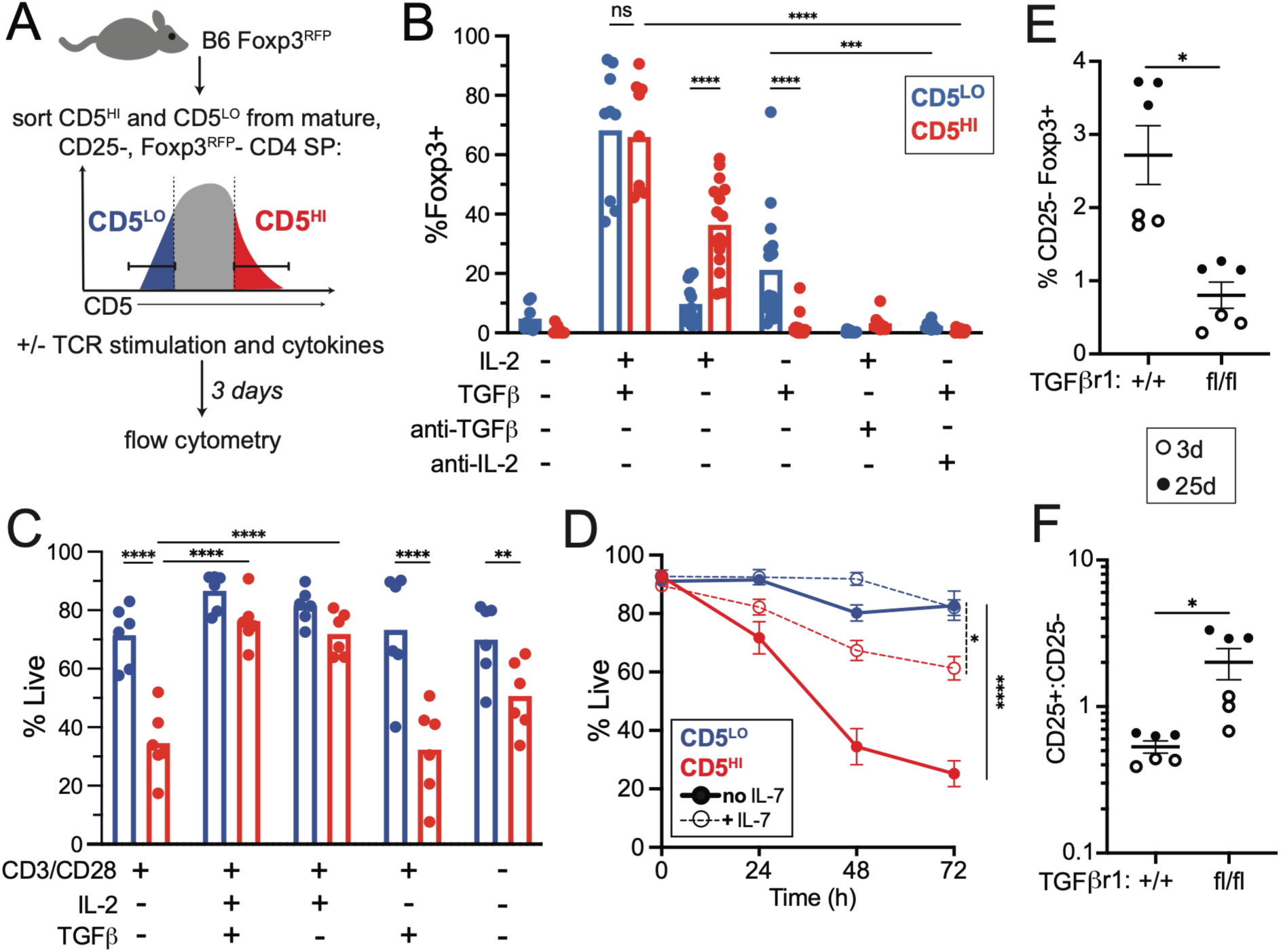
Functional tuning during positive selection correlates with differential responses to Treg inducing cytokines. A. Experimental schematic for data in panels C, D. Thymocytes from Foxp3^RFP reporter mice were sorted for mature CD4 SP (CD8-CD4+CD69-CD24^LO^) and for the top and bottom 25% of CD5 (designated CD5^HI^ or CD5^LO^). Sorted thymocytes were cultured with plate-bound anti-CD3/CD28 antibodies and the indicated cytokines for 72 hours and analyzed by flow cytometry. B. Plots show the % of Foxp3^RFP reporter+ cells out of live CD4 SP. Where indicated, exogenous IL-2 and/or TGFβ or anti-IL-2 or anti-TGFβ blocking antibodies were added to cultures. Each dot represents one replicate from at least five experiment. C. Same samples from B showing the % of live cells out of the CD4 SP gate for each condition. Samples cultured for 72 hours without anti-CD3/CD28 stimulation are shown for comparison. D. Thymocytes from B6 WT mice were cultured in the presence or absence of IL-7 for the indicated times and analyzed by flow cytometry. Plot shows the % of live cells out of the mature (CD8-CD4+CD69-CD24^LO^) CD4 SP gate for the top and bottom 25% of CD5 (CD5^HI^ or CD5^LO^). Thin dotted lines represent no added IL-7 and thick solid lines represent 20 ng/mL added IL-7. Each dot represents one replicate from a total of four experiments. E,F. CD25+ and CD25- thymic Treg populations from TGFβr1^fl/fl^ x CD4^Cre^ mice (control animals were TGFβr1^+/+^ x CD4^Cre^) were analyzed. Mice age 3d (open circles) and 25d (closed circles) were used. D. Ratio of CD25+ to CD25- Tregs in each condition. E. % CD25+ Foxp3+ and CD25- Foxp3+ of CD4 SP thymocytes for each condition.

It has been suggested that cytokines promote Treg development by enhancing survival as well as directly inducing Foxp3 gene expression (Ouyang *et al*., 2010; Tai *et al*., 2013; Hu *et al*., 2017; Dikiy *et al*., 2021). We therefore assessed survival of CD5^HI^ and CD5^LO^ CD4 SPs after 72h of TCR stimulation in Treg induction cultures. We observed that CD5^LO^ CD4 SPs survived better under most conditions compared to CD5^HI^ CD4 SPs **(Fig. 4C)**. Moreover, added IL-2 substantially improved the survival of CD5^HI^, but not CD5^LO^, CD4 SP. We noted that the survival advantage of CD5^LO^ CD4 SPs was observed even in the absence of TCR stimulation. To extend this observation, we cultured WT thymocytes with or without the survival-promoting cytokine IL-7 and assessed the % of live cells within the CD5^HI^ and CD5^LO^ CD4 SP gates at different time points. We observed that mature CD5^LO^ CD4 SP showed better survival than CD5^HI^ CD4 SP, especially in the absence of IL-7 (**Fig. 4D**). This is in line with previous observations of mature CD5^LO^ and CD5^HI^ CD8 T cells (Palmer *et al*., 2011). Taken together these results imply that positive selection imprints CD4 SP thymocytes with distinct potential to survive and respond to Treg-inducing cytokines; differences that may underlie their tendency to give rise to CD25+ or CD25- Tregs upon agonist selection.

### Loss of TGFβ signaling has a greater impact on CD25- compared to CD25+ thymic Tregs

Given that TGFβ potently induces Treg development from CD5^LO^ CD4 SP thymocytes *in vitro* (**Fig. 4B**), and that TCR transgenic CD5^LO^ CD4 SP develop into CD25- Tregs in thymic slice cultures (**Fig. 1**) we hypothesized that thymic CD25- Tregs would be more dependent on TGFβ for their development *in vivo*. Previous studies demonstrated that loss of TGFβ signaling due to a T cell specific gene ko of *Tgfβr1* led to overall reduced Treg development in neonatal mice (Liu *et al*., 2008), however the specific impact on CD25- versus CD25+ thymic Tregs was not examined. To address this question, we examined CD25 expression on thymic Tregs from mice with a T cell-specific TGFβ receptor 1 knockout (*Tgfβr1^fl^*^/fl^ crossed to CD4^Cre^). Consistent with previous studies, ko of *Tgfβr1* led to an overall loss of thymic Tregs at 3 days, while IL-2-driven expansion led to a recovery of Treg numbers by day 25 (Liu *et al*., 2008). However, at both time points *Tgfβr1^fl^*^/fl^ mice displayed a significant decrease in thymic CD25- Tregs and a higher ratio of CD25+ to CD25- Tregs than their *Tgfβr1*^+/+^ litermates (**Fig. 4E, F, Suppl. Fig 5A**). These results indicate that TGFβ plays a more important role in the thymic development of CD25- Tregs compared to CD25+ Tregs.

### CD25- thymic Tregs remain CD25- as peripheral central Tregs

Although it has been proposed that CD25-Foxp3+ thymocytes are precursors to CD25+ thymic Tregs (Marshall *et al*., 2014; Owen *et al*., 2019), our evidence that CD25- thymic Tregs derive from distinct subsets of CD4 SP and exhibit distinct cytokine requirements led us to hypothesize that they instead represent a separate lineage of thymic derived Tregs that lacks constitutive CD25 expression. To test this hypothesis, we generated Tregs *in vivo* from TCR transgenic models to ask whether clonal Tregs retained their thymic CD25 expression patterns after leaving the thymus. For the BDC2.5 TCR, we extended an earlier study (Lin *et al*., 2022) in which thymic Treg induction in neonatal BDC2.5 TCR transgenic mice was driven by injection of the agonist 2.5HIP peptide conjugated to an anti-Langerin antibody to target thymic DC (Ab-2.5HIP). While a large population of thymic CD25- Tregs was reported, the peripheral Treg populations were not characterized in this study. To address this question, we injected one day old neonatal BDC2.5 mice i.p. with Ab-2.5HIP and analyzed thymi, spleens, and mesenteric lymph nodes for Foxp3 and CD25 expression several days later. Consistent with the previous study (Lin *et al*., 2022) and our thymic slice experiments (**Fig. 1**), a substantial population of CD25- Tregs was observed in the thymus of Ab-2.5HIP injected, but not control mice (**Fig. 5A-C**). Moreover, in the spleen and mesenteric lymph nodes of BDC2.5 neonates, we also observed predominantly CD25- Tregs (**Fig. 5B, C**). These peripheral CD25-Tregs expressed Helios (**Suppl. Fig 6A**) implying that they were thymically derived Tregs (Thornton *et al*., 2010). These results indicate that, in the presence of agonist ligand, thymocytes expressing the BDC2.5 TCR generated thymic CD25- Tregs that remained CD25- after they migrated to the periphery.

**Figure 5.**
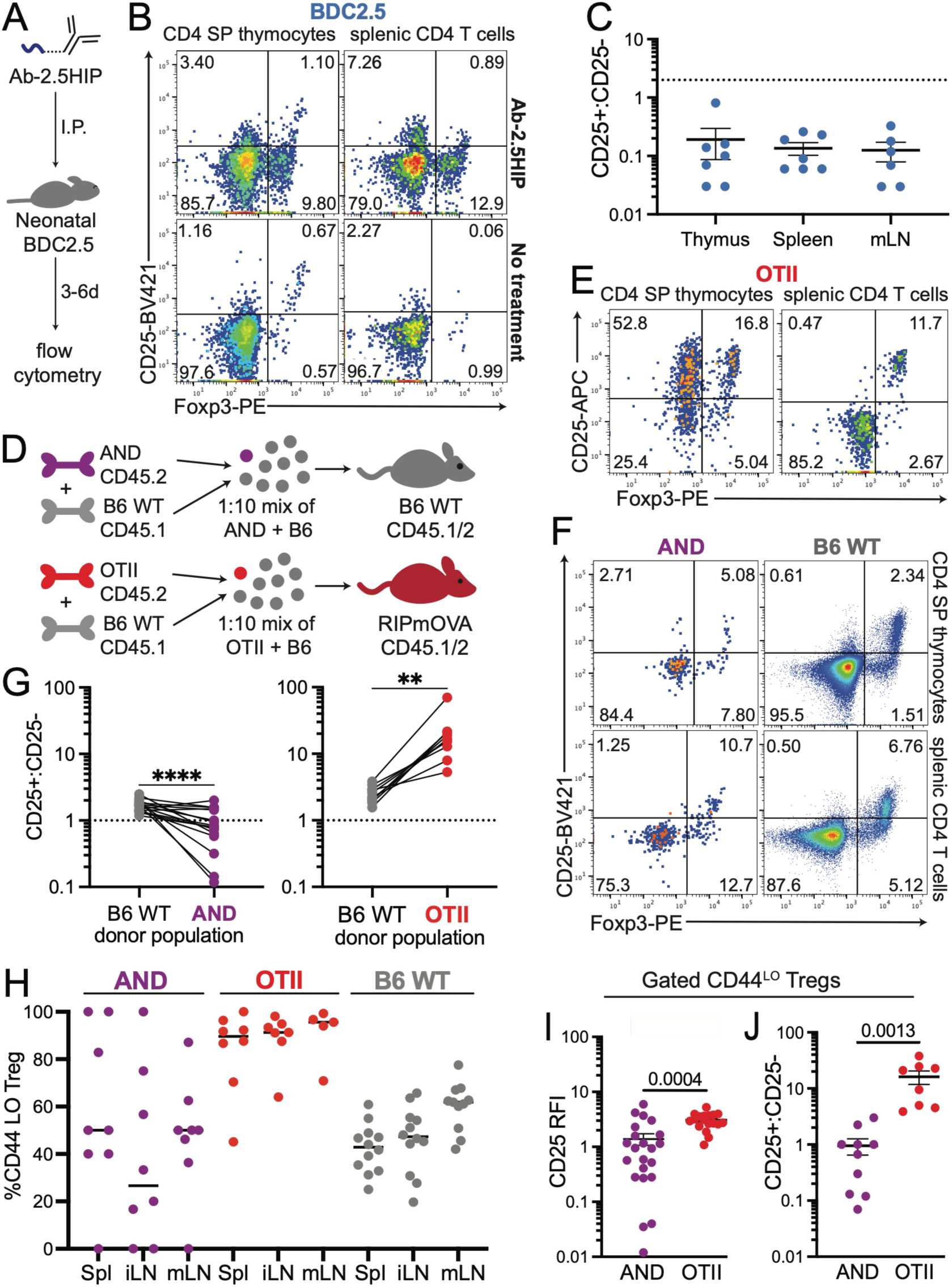
CD25- thymic Tregs remain CD25- as peripheral central Tregs. A. Schematic for neonatal BDC2.5 Ab-2.5HIP I.P. injections. 2.5 µg Ab-2.5HIP was I.P. injected into 1d old BDC2.5 neonates. 3-6d later, thymi, spleens, and mesenteric lymph nodes (mLN) were taken for flow cytometry. B. Representative flow plots of CD25 and Foxp3 expression on gated BDC2.5 CD4 SP thymocytes (right) or CD4 T cells (left) either injected with 2.5 µg Ab-2.5HIP (top) or untreated (bottom). C. Ratios of CD25+ to CD25- Foxp3+ Tregs from neonatal BDC2.5 CD4 SP thymocytes and CD4 T cells from spleen and mLN. D. Schematic for low frequency TCR transgenic bone marrow chimeras. Top: RIPmOVA hosts were irradiated and then reconstituted with a 1:10 mixture of congenically distinct OTII (CD45.2) and B6 WT (CD45.1) bone marrow. Bottom: B6 WT hosts were irradiated and then reconstituted with a 1:10 mixture of congenically distinct AND (CD45.2) and B6 WT (CD45.1) bone marrow. E-J. Thymus, spleen, and lymph nodes from chimeric mice were analyzed by flow cytometry. Representative flow plots of CD25 and Foxp3 expression on gated donor-derived CD4 SP thymocytes (left) and splenic CD4 T cells (right) from (E) OTII, or (F) AND (top) and B6 WT (bottom) donor populations. G. Compiled data from pooled lymph nodes and spleen of chimeric mice. showing the ratios of CD25+Foxp3+ to CD25-Foxp3+ Tregs from donor-derived AND (left) or OTII (right) and B6 WT CD4 T cells. Black lines indicate paired comparisons of TCR transgenic and B6 WT CD4 T cells from the same samples. H. Compiled data showing the % of CD44^LO^ cells out of Foxp3+ CD4+ T cells from spleen (Spl), inguinal LN (iLN), and mesenteric LN (mLN) for the indicated donor populations. I. Compiled data showing relative expression (gMFI) of CD25 on AND and OTII CD44^LO^ Foxp3+ Tregs normalized to the gMFI of B6 WT CD44^LO^ Foxp3+ Tregs from the same chimera. J. Compiled data showing the ratio of CD25+ CD44^LO^ Foxp3+ to CD25- CD44^LO^ Foxp3+ Tregs from AND and OTII CD4 T cells from spleen and lymph nodes of mixed bone marrow chimeras. Compiled from five experiments of 2-3 mice each for AND and three experiments of 2-3 mice each for OTII and BDC2.5. Horizontal lines represent the mean (and vertical lines represent +/- SEM. For statistical analysis, paired t-test was used. **=P<0.01, ***=P<0.001.

To explore the peripheral phenotypes of AND and OTII TCR transgenic Tregs, we generated low frequency hematopoietic chimeras by reconstituting irradiated hosts with TCR transgenic bone marrow mixed with an excess of WT donor bone marrow **(Fig. 5D).** For OTII, irradiated hosts were RIPmOVA transgenic mice in which the model antigen OVA is expressed by a subset of medullary thymic epithelial cells and islet cells of the pancreas (Kurts *et al*., 1996). For AND, we used B6 WT hosts, taking advantage of the endogenous agonist ligand **(Suppl. Fig. 2)**. We analyzed thymi from chimeric mice for CD25 and Foxp3 expression and observed that, similarly to thymic slice experiments **(Fig. 1)**, OTII CD4 SP thymocytes developed into predominantly CD25+ Tregs, while AND CD4 SP thymocytes developed into a mix of CD25+ and CD25- Tregs **(Fig. 5E,F)**. In the peripheral Treg compartment, AND Tregs had a lower ratio of CD25+ to CD25- Tregs compared to WT donor-derived Tregs in the same chimeric mice, while the opposite was true for OTII Tregs **(Fig. 5G**). This provides further evidence that individual TCRs confer a preference for either a CD25- or CD25+ Treg fate that begins in the thymus and is maintained after they leave the thymus and migrate to peripheral lymphoid tissues.

To further investigate the phenotype of AND and OTII peripheral Tregs, we used CD44 expression to distinguish “central” Tregs (CD44^LO^) that phenotypically resemble newly developed thymic Tregs, versus “effector” Tregs (CD44^HI^) that have undergone further activation in the periphery (Smigiel *et al*., 2014). While OTII Tregs were predominantly CD44^LO^, AND Tregs showed considerable variation in CD44 expression **(Fig. 5H)**. Focusing on the CD44^LO^ peripheral T cell population further confirmed the skewing of CD25 expression between AND and OTII peripheral Tregs. On average, AND CD44^LO^ Tregs displayed lower CD25 expression and a reduced ratio of CD25+ to CD25- Tregs compared to OTII CD44^LO^ Tregs **(Fig. 5I,J).** Interestingly, while the expression of CD25 and CD44 on WT and OTII Tregs showed an inverse correlation **(Suppl. Fig. 6B,C)**, consistent with previous findings (Smigiel *et al*., 2014), AND Tregs showed a positive correlation between these markers **(Suppl. Fig. 6D**). These results suggest that thymic CD25- Tregs remain CD25- as CD44^LO^ peripheral Tregs but have the potential to upregulate CD25 in response to further activation signals in the periphery.

### CD25- thymic Tregs and CD25- splenic central Tregs share a similar TCR repertoire

To extend these observations beyond TCR transgenic models, we compared the TCR repertoire of CD25+ and CD25- thymic and peripheral Tregs from fixed TCRβ transgenic mice. These mice exhibit relatively normal immune responses, but their TCR diversity is limited to the TCRα chain, enabling quantitative repertoire analyses by sequencing of TCRα genes from different T cell populations (Hsieh *et al*., 2006; Pacholczyk *et al*., 2006). We crossed RT3 TCRβ transgenic mice (Malchow *et al*., 2016) with Foxp3^RFP^ reporter mice to enable FACS isolation of Tregs, and TCRα-/- mice to ensure that each T cell expressed only a single TCRα chain. We then isolated CD25+ and CD25- thymic Tregs (Foxp3^RFP^+CD4+CD8-) cells and splenic central Tregs (Foxp3^RFP^+CD4+CD8-CD44^LO^) from 2-3 individual mice, performed TCRα gene sequencing, and compared the repertoires across the samples using the Morisita-Horn (MH) similarity index (**Fig. 6A**). The TCRα repertoire of CD25- Tregs was strikingly similar between thymus and spleen CD25- cTreg samples (MH index 0.26-0.43), while showing very little similarity to CD25+ Tregs (MH index <0.06) (**Fig. 6B,C**). In addition, the TCRα repertoire of thymic CD25+ Tregs was more similar to CD25+ splenic cTregs than to CD25- Tregs. It also is noteworthy that the mouse-to-mouse repertoire similarity between CD25- Treg samples (MH index 0.26-0.63) was greater than for CD25+ Tregs (MH index 0.06-0.22) (**Fig. 6B, C**) indicating that CD25-Tregs have a more stable and convergent TCR repertoire.

**Figure 6.**
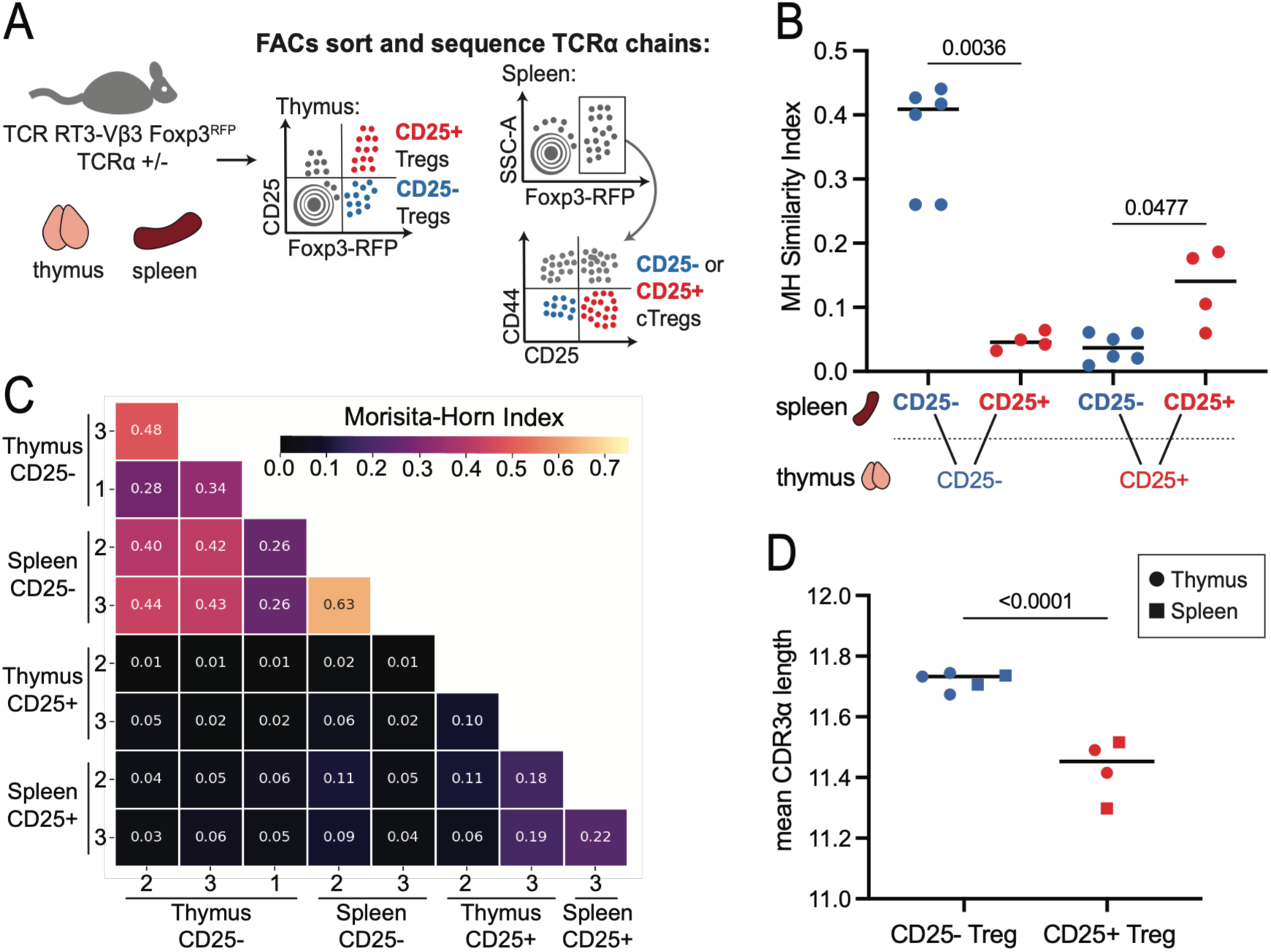
CD25- thymic Tregs and CD25- splenic central Tregs share a similar TCR repertoire. A. Experimental schematic for TCR repertoire analyses of CD25- and CD25+ Tregs from limited TCR diversity mice. Thymic CD25+ or - Tregs and splenic CD25+ or - cTreg (CD44^LO^) populations were sorted from three RT3 TCRβ transgenic Foxp3^RFP^ TCRα-/+ mice and subjected to TCRα sequence analyses. B-C. Morisita-Horn similarity index scores for pairwise comparisons between thymic CD25- Treg populations compared to splenic CD25+ or - cTregs (B), or for all samples (C) are shown. In (B) each dot corresponds to a pairwise comparison between thymus and spleen samples from individual mice. D. Comparison of average CDR3α length between CD25+ and - Tregs. Circles indicate thymic and squares indicate splenic Tregs. B,D. Average values are indicated by horizontal bars. For statistical analysis, (B) unpaired t-test and (D) one way analysis of variance (ANOVA) with multiple comparisons was used.

Focusing on the most prevalent recurrent clones, we observed that, while many CD25- thymic Treg clones were present at similar frequencies in the CD25- splenic cTreg repertoire, very few were present in either the thymic or splenic CD25+ Treg repertoires (**Suppl. Fig 7A**). Likewise, the most prevalent recurrent clones in the CD25+ thymic Treg repertoire were present in the CD25+ splenic cTreg repertoire at higher frequencies than either the thymic or splenic CD25- Treg repertoires (**Suppl. Fig 7B**). We also noted that CD25+ Tregs had shorter CDR3αs on average than CD25- Tregs (**Fig 6D, Suppl Fig. 7C**). Given previous reports that CD5^HI^ CD4 SPs have shorter CDR3s than CD5^LO^ CD4 SPs (Stadinski *et al*., 2016; Textor *et al*., 2023), these data provide additional support for the notion that thymic CD25- Tregs are derived from CD5^LO^ CD4 SP thymocytes. Altogether, these TCR repertoire analyses further support the hypothesis that CD25- and CD25+ thymic Tregs are distinct lineages that maintain their CD25 expression pattern in the periphery.

## Discussion

While it is well established that functional tuning during positive selection impacts how naïve T cells respond to antigen and cytokine stimulation, its impact on thymic Treg development has remained unexplored. Here we provide evidence that the strength of the TCR signal experienced by CD4 SP thymocytes during positive selection in the thymic cortex predisposes them to adopt either a classic CD25+ or an alternative CD25- Treg fate (Treg^alt^) upon encounter with agonist ligands in the medulla (**Suppl. Fig. 8A**). Mechanistically, we provide evidence that the level of self-reactivity experienced during positive selection impacts responses to Treg-inducing cytokines, such that CD4 SP with low cortical self-reactivity (CD5^LO^) are more dependent on TGFβ, whereas CD4 SP with moderate cortical self-reactivity (CD5^HI^) are more dependent on IL-2. As a result, loss of TGFβ signaling in thymocytes *in vivo* leads to a selective loss of CD25- Tregs. We also provide evidence that CD25- thymic Tregs tend to maintain their CD25- phenotype as peripheral central Tregs and harbor a distinct, stable TCR repertoire. These data provide a new perspective on how the thymic selection of Tregs shapes peripheral Treg heterogeneity.

While constitutive CD25 expression is a classic hallmark of Tregs, not all peripheral Foxp3+ CD4 T cells express high levels of CD25 (Zelenay *et al*., 2005; Ono *et al*., 2006; Coleman *et al*., 2012; Smigiel *et al*., 2014; Wyss *et al*., 2016). Moreover, conventional (Foxp3-) T cells transiently upregulate CD25 upon stimulation, further complicating its use as a cell type marker. We propose that alternative Tregs lack constitutive CD25 expression but can transiently upregulate CD25 upon stimulation (**Suppl. Fig. 8B, dashed arrow**). While the ability of thymic CD25-Foxp3+ CD4 SP thymocytes from non-TCR transgenic mice to give rise to some CD25+ Tregs upon IL-2 exposure or *in vivo* transfer (Marshall *et al*., 2014; Owen *et al*., 2019) has been taken as evidence that they represent precursors to thymic CD25+ Tregs, it is unclear whether CD25 expression on the recovered population was transient or constitutive. Moreover, given the polyclonal nature of the transferred population, the low cell recovery, and the fact that roughly equal numbers of CD25+ and CD25- Tregs were detected in the recovered population, these data are fully consistent with our interpretation that CD25- thymic Tregs can give rise to an alternative lineage of CD25- peripheral Tregs. Future studies of alternative Tregs will be aided by tracking of clonal Treg^alt^ cells, as well as approaches that distinguish between constitutive vs. induced CD25 expression.

Our data shed light on the relative roles of IL-2 and TGFβ during thymic Treg development. Specifically, our data confirm that both IL-2 and TGFβ are needed for optimal Treg induction *in vitro* from thymic CD4 SP, but point to differential cytokine dependence for CD5^HI^ versus CD5^LO^ CD4 SP. CD5^LO^ CD4 SP show a greater dependence on TGFβ for Foxp3 induction, while CD5^HI^ CD4 SP are more dependent on IL-2. Moreover, our *in vivo* data showing that loss of TGFβ signaling in thymocytes causes a greater loss of CD25- compared to CD25+ thymic Tregs, together with prior *in vivo* data showing that CD25+ Tregs are more dependent on IL-2 than CD25- Tregs (Marshall *et al*., 2014; Apert *et al*., 2022) align with the *in vitro* data, and support the view that TFGβ is the more important inducer of CD25-Treg development from CD5^LO^ CD4SP, while IL-2 is more important for driving CD25+Tregs from CD5^HI^ CD4 SP. While there is evidence that IL-2 can prevent the death of CD4 SP experiencing strong TCR signals (Tai *et al*., 2013; Hu *et al*., 2017), our data indicate that added IL-2 promotes the survival of CD5^HI^, but not CD5^LO^ CD4 SP (**Fig. 4E**). The observation that CD5^LO^ CD4 SP generally exhibit superior survival *in vitro* may help to explain their reduced dependence on IL-2. We speculate that, for CD5^LO^ CD4SP, a relatively low STAT5 signal provided by trace amounts of IL-2 or other common gamma chain cytokines such as IL-15, together with TGFβ, provide optimal signals to drive their development into Treg^alt^ cells.

In addition to exhibiting a greater dependence on IL-2 for Treg development, CD5^HI^ CD4 SP also produce more IL-2 immediately following TCR stimulation, as reported previously (Persaud *et al*., 2014; Zinzow-Kramer, Weiss and Au-Yeung, 2019) and confirmed here. It is interesting to consider the possible mechanistic connection between IL-2 production by CD5^HI^ CD4 SP and their tendency to develop as CD25+ Tregs remains unclear. While an initial burst of IL-2 may act in an autocrine fashion to promote CD25+ Treg development (Hemmers *et al*., 2019), this would likely be a transient event, given that Foxp3 potently suppresses IL-2 expression. Another non-mutually exclusive possibility is that CD5^HI^ autoreactive CD4 SP provide IL-2 in a paracrine fashion to CD5^HI^ Treg precursors of related specificity that interact with the same APC. Given the rarity of thymic APC presenting particular tissue restricted antigens (Derbinski *et al*., 2008), and the efficient scanning by medullary thymocytes (Le Borgne *et al*., 2009), such multi-cell interactions between an APCs and two or more CD4 SP thymocytes reactive with the same tissue-restricted antigen may occur relatively frequently. Co-localization of IL-2-producing CD4 T cells and Treg precursors could promote a balanced production of autoreactive CD4 T cells and Tregs with related specificity (Klein, Robey and Hsieh, 2019; Klawon *et al*., 2025). This would be in line with the "Buddy Hypothesis": which posits that autoreactive T cells are released from the thymus along with a Treg buddy of related specificity to keep them in check (Hsieh, Lee and Lio, 2012). It is also interesting to consider that the strong IL-2 dependence of CD5^HI^-derived Tregs is likely retained after they leave the thymus, making them well poised to limit IL-2 availability to their autoreactive conventional T cell "buddies" in SLO. On the other hand, the intrinsic survival advantage of CD5^LO^ CD4 SPs may be retained after their conversion to Treg^alt^ cells, thus reducing their dependence on IL-2 and better allowing them to survive in tissues.

Altogether, our study points to a role for positive selection tuning in thymic Treg development and provides evidence for a previously unappreciated alternative Treg lineage. Although the current study focused on the thymic development of Treg^alt^ cells, the lack of constitutive CD25 expression and distinct functional tuning they received during positive selection suggests that Treg^alt^ cells may occupy distinct anatomical and functional peripheral niches compared to classic Tregs. Given the intense interest in harnessing Tregs for therapeutic purposes (Wang *et al*., 2025), a full understanding of Treg^alt^ cells is a high priority for future studies.

## Materials and Methods

### Mice and bone marrow chimeras

The following mice were obtained from the Jackson Laboratory and were bred and maintained under specific pathogen-free (SPF) conditions in accordance with guidelines approved by the Institutional Animal Care and Use Committees at the University of California, Berkeley: C57BL/6 (B6, #000664), C57BL/6 CD45.1 (B6 CD45.1, #002014), B10.BR (#004804), NOD/ShiLtJ (NOD, #001976), BDC2.5 TCR (#004460, Katz 1993), RIP-mOVA (#005431, Kurts 1996), RAG1-/- (#002216), RAG2-/- (#008449), TCRα-/- (#002116), AND TCR (#002408, Kaye 1989), and OTII TCR (#004194, Barnden 1998). AND and OTII TCR transgenic mice were crossed onto RAG1-/- and RAG2-/- backgrounds, respectively. *Tgfbr1*^f/f^ mice (#028701) and CD4-Cre mice (#022071) were both obtained from the Jackson Laboratory, and *Tgfbr1*^f/f^CD4-Cre mice were bred under SPF conditions in accordance with guidelines approved by the Animal Care and Use Committees of the National Institute of Dental and Craniofacial Research (NIDCR). Foxp3-RFP (#008734, Jackson Laboratory) mice were generously provided by the DuPage lab. RT3-TCRβ transgenic mice (Malchow et al, 2016) were generously provided by the Savage Lab and were crossed to Foxp3^RFP reporter and TCRα KO in our facility. C57BL/6 CD45.1 mice were crossed with B10.BR mice and resulting F_1_ progeny (I-A^b^/I-E^k^) were used for thymic slices and BMDCs where MCCp presentation on I-E^k^ was required. Low frequency TCR transgenic bone marrow chimeras were generated by transferring a 1:10 mixture of TCR transgenic and non-transgenic B6 CD45.1 (5 x 10^6^ bone marrow cells total) intravenously into lethally irradiated 4-to 6-week old B6 or RIPmOVA CD45.1/CD45.2 hosts. Chimeras were allowed to reconstitute for 5-6 weeks before analysis.

### Ab-2.5HIP conjugation and intraperitoneal (I.P.) injection

Ab-2.5HIP was assembled as previously described (Lin *et. al.* 2022). Briefly, the 2.5HIP peptide with an N-terminal cysteine (LQTLALWSRMD-C) was purchased from GenScript. The anti-Langerin antibody with a dockerin domain as well as the cohesin-thiopyridine was kindly provided by the Zurawski Laboratory (Baylor Institute for Immunology research). To produce Ab-2.5HIP, cohesin and 2.5HIP were first incubated together overnight at room temperature (RT) to produce a cohesin-2.5HIP conjugate. The anti-Langerin-dockerin antibody was then mixed with the cohesin-2.5HIP conjugate and PBS containing calcium, and incubated for 30 minutes at RT. The mixture was then passed through a 7K MWCO Zeba-Spin Column (ThermoFisher #89877) to remove any unconjugated protein. For neonatal IP injections, 2.5 µg of Ab-2.5HIP was I.P. injected into one day old BDC2.5 neonates.

### Thymus, spleen and lymph node isolation

Spleens, lymph nodes, and/or thymi were dissected from euthanized mice and mechanically dissociated in homogenizers containing complete RPMI (cRPMI: 10% FBS, 8 mM HEPES, 1 mM sodium pyruvate, 55 μM β-mercaptoethanol, non-essential amino acids, L-glutamine, and penicillin-streptomycin). Homogenized tissues were pipetted through 70 μm filters to produce single cell suspensions. Samples were centrifuged and resuspended in ACK lysis buffer to remove red blood cells. After 3-5 minutes ACK lysis was quenched by the addition of 5 mL of cRPMI. Cells were centrifuged again and resuspended at appropriate concentrations for downstream analyses.

### Bone marrow dendritic cell generation and experimental use

Bone marrow was harvested from the femurs and tibias of mice and red blood cells were lysed with ACK buffer. Cells were resuspended at a concentration of 0.5 x 10^6^/mL in complete DMEM (cDMEM: 10% FBS, 55 μM β-mercaptoethanol, 4.5g/L glucose, L-glutamine, non-essential amino acids, penicillin/streptomycin) supplemented with 10 ng/mL GM-CSF (Peprotech #315-03) and 10 ng/mL IL-4 (Peprotech #214-14) and plated in 100mm petri dishes (Corning #353003). A half media change was performed at day 2 and a full media change was performed at day 5. At day 7, media was removed and cold 2 mM EDTA in PBS was added for 10 minutes to dissociate loosely adherent cells. For thymic slice experiments, BMDCs were resuspended at 1 x 10^6^ cells/mL in pre-warmed 1X HBSS with 1 μM agonist peptide and incubated at 37°C for 30 minutes. Peptide-loaded BMDCs were washed 3x with a 5- to 10-fold excess of 1X HBSS to remove unbound extra peptide before being resuspended in 10 μL of cRPMI and overlaid onto thymic slices 3-4 hours after overlay of TCR transgenic thymocytes. BMDCs were allowed to migrate into slices for 2.5-3 hours before excess cells were gently washed off. For *in vitro* stimulations, 2.5 x 10^5^ BMDCs were plated in 48-well plates containing 500 μL cRPMI with 1 μM agonist peptide and 5 x 10^5^ TCR transgenic thymocytes.

### Generation of thymic tissue slices

Thymic slices were prepared as previously described (Ross 2016, McIntyre 2023). Briefly, thymic lobes were dissected from mice, embedded in 4% low melt temp agarose and sliced on a vibratome into 400-500 μm sections. Slices were cultured on 0.4 um transwell cell culture inserts (Corning #353090) placed in cRPMI. 0.5-2x10^6^ TCR transgenic thymocytes were resuspended into 10 μL cRPMI, overlaid onto the slice, and were allowed to migrate into the tissue for 3-4 hours, after which excess thymocytes were washed off. To distinguish AND or OTII TCR transgenic from slice resident thymocytes, congenically distinct TCR transgenic thymocytes (CD45.2+) and thymic slices (CD45.1+) were used. For experiments where overlaid thymocytes were congenically identical to slices (such as BDC2.5 thymocytes on NOD slices), overlaid TCR transgenic thymocytes were labeled with CFSE or CellTrace Violet before use. For “shared niche” and “separate niche” slice experiments where AND and OTII TCR transgenic thymocytes were present at a 1:1 ratio in the same slice, AND and OTII thymocytes were mixed at a 1:4 ratio (a mixture of 0.5 x 10^6^ AND thymocytes + 2 x 10^6^ OTII thymocytes were overlaid per slice) before overlay to account for different migratory capacities. AND thymocytes were labeled with CellTrace Violet (CTV) before mixing with OTII thymocytes so that AND thymocytes (CTV+CD45.2+) and OTII thymocytes (CTV-CD45.2+) could each be distinguished from slice resident thymocytes (CD45.1+/CD45.2+).

### Flow cytometry

Lymphocytes were stained using the following antibodies: CD4 (clone RM4-5 BD Biosciences) CD8α (clone 53-6.7 BD Biosciences), CD5 (clone 53-7.3 BD Biosciences), CD24 (clone M1/69, BioLegend), CD69 (clone H1.2F3, BioLegend), CD25 (clone PC61.5, eBioscience), CD44 (clone IM7, BD Biosciences), CD45.1 (clone A20, Tonbo Biosciences) CD45.2 (clone 104, BD Biosciences), TCR Vα2 (clone B20.1, BioLegend), TCR Vα11 (clone RR8-1, eBioscience) Foxp3 (FJK-16s, eBioscience), IL-2 (JES6-5H4, BioLegend). Cells were first stained with Zombie NIR or Yellow Viability Dye (NIR: BioLegend #423105, Yellow: #423103) for 20 minutes at 4°C, then washed with 1X PBS and stained with antibodies against surface proteins in 2.4G2 supernatant for 30 minutes at 4°C before washing and resuspension in FACs buffer (1X HBSS with 1% BSA and 0.1% sodium azide). For intracellular Foxp3 stains, cells were fixed and permeabilized using the Foxp3/Transcription Factor Staining Buffer Set (eBioscience 00-5523-00) according to manufacturer protocol. For intracellular IL-2 stains, cells were pre-treated with 1X Protein Transport Inhibitor Cocktail (eBioscience 00-4980-93) for at least 4 hours before staining. Viability dye and surface proteins were stained as described above, and intracellular IL-2 was stained after fixation and permeabilization using the Cytofix/Cytoperm Fixation/Permeabilization Kit (BD Biosciences #554714) according to manufacturer protocol. Data acquisition was performed on a BD Symphony A3 or BD LSR Fortessa X-20 and data analysis was performed with FlowJo V10. For live cell sorting, B6 Foxp3^RFP double positive and CD8 single positive thymocytes were depleted using a biotinylated CD8α antibody (clone 53-6.7, eBioscience) and the Mouse Biotin Positive Selection Kit II (StemCell Technologies #17665) before viability and surface staining. Populations from enriched CD4 SP thymocytes were then sorted on a BD FacsAria Fusion.

### PMA and Ionomycin *in vitro* stimulation

For *in vitro* stimulations using phorbol-12-myristate-13-acetate (PMA) and ionomycin, TCR transgenic thymocytes were resuspended in cRPMI with 25 ng/mL PMA, 0.1 μg/mL ionomycin, and 1X Protein Transport Inhibitor Cocktail. Thymocytes were plated in 48-well format at 0.5 x 10^6^ cells/well, and stim/no stim conditions were performed in triplicate. Thymocytes were incubated at 37°C for 16 hours before cells were stained for analysis.

### *In vitro* Treg development

For *in vitro* Treg development assays, thymocytes from B6 Foxp3^RFP mice were sorted by top and bottom 20% CD5 expressing (CD5^HI^ and CD5^LO^, respectively) mature (CD24^LO^ CD69-) conventional (CD25-Foxp3-) CD4 SP (CD4+CD8-). Sorted CD4 SP were resuspended in cRPMI with one or more of the following: 50 U/mL IL-2 (Peprotech #200-02), 5 ng/mL TGFβ (Peprotech #100-21), 10 μg/mL anti-IL-2 (clone JES6-5H4, InVivoMab #BE0042), and/or 0.5 μg/mL anti-TGFβ (clone 1D11, R&D Systems #MAB1835), and plated at 1 x 10^5^ cells/mL in 96-well flat bottom plates. To provide TCR stimulation for CD4 SP thymocytes, plates were pre-coated with 1X PBS containing 1 μg/mL each of anti-CD3 (clone 145-2C11, Tonbo Biosciences) and anti-CD28 (clone 37.51, Tonbo Biosciences) overnight at 4°C. PBS with anti-CD3/28 was aspirated from wells before cells were plated. Cells were incubated for 72h at 37°C before analysis. For experiments assessing survival of B6 WT thymocytes with and without IL-7, whole thymocyte single cell suspensions were resuspended at 2 x 10^5^ cells/mL in cRPMI with or without 20 ng/mL IL-7 (Peprotech #217-17), plated in 96-well plates and cultured at 37°C for up to 72 hours before analysis.

### TCRα sequencing

Single cell suspensions were prepared from thymi and spleens from RT3-TCRβ transgenic Foxp3^RFP TCRα+/- mice. Thymocytes were enriched for CD4 T cells using an anti-CD8 biotin antibody (Invitrogen #13-0081-85), and splenocytes were enriched for CD4 T cells using anti-CD8 and anti-CD19 (Invitrogen #13-0199-82) biotin antibodies. Lymphocyte fractions were depleted of unwanted biotin-bound cells using the EasySep Mouse Streptavidin RapidSpheres isolation kit and protocol (STEMCELL #19860). Enriched CD4 T cell fractions were stained for CD4, CD8, CD25 and CD44 (for staining spleen samples only) in 2.4G2 supernatant on ice for 30 minutes, washed with FACs buffer, and resuspended in OptiMEM + 2% FBS before sorting on an Aria Fusion cell sorter. 10,000-15,000 CD25+ or CD25- thymic Tregs (Foxp3^RFP+CD4+CD8-) and splenic Tregs (Foxp3^RFP+CD4+CD8-CD44^LO^) were recovered. Sorted cells were washed with 1X PBS before cell lysis and RNA extraction was performed according to manufacturer protocol using the RNAqueous Micro Kit (Invitrogen #AM193110). Purified RNA was used for library preparation using the RNA-iR-Complete Dual Index Primer Kit from iRepertoire. After successful library generation, libraries were pooled at equimolar ratios and sequenced using the Illumina MiSeq i100 platform at the UC Berkeley genomics sequencing laboratory. Initial analyses of TCRα sequence data was performed using iRepertoire Standard Data Analysis package, and Morisita-Horn similarity indexes were determined using Python.

## Acknowledgements

We thank Silvia Ariotti for initiating the project, Karly Ortega and Sophie Keshishian for technical assistance, Pete Savage for providing the RT3 TCR transgenic mice, and Tom Mann and Michel DuPage for comments on the manuscript. We thank Sandy Zurawski for providing anti-Langerin-dockerin antibody and cohesin-thiopyridine protein, and Matt Bettini for providing protocols and advice for regarding Ab-2.5HIP conjugation and neonatal administration. We thank the Cancer Research Lab Flow Cytometry Core Facilities at UC Berkeley, including Kartoosh Heydari, Melaine Delcroix, and Harman Dhaliwal for their help operating the flow cytometers and cell sorters. I.B., Y. K., and E.R. were funded by NIH NIAID RO1AI064227. D.Z., and WJ.C. were supported by the Intramural Research Programs of NIDCR, National Institute of health (NIH), USA.

