## Supplementary Figures for "TCR signal strength during positive selection shapes CD25 expression patterns on thymically derived regulatory T cells"

**Supplementary Figures and Figure Legends**

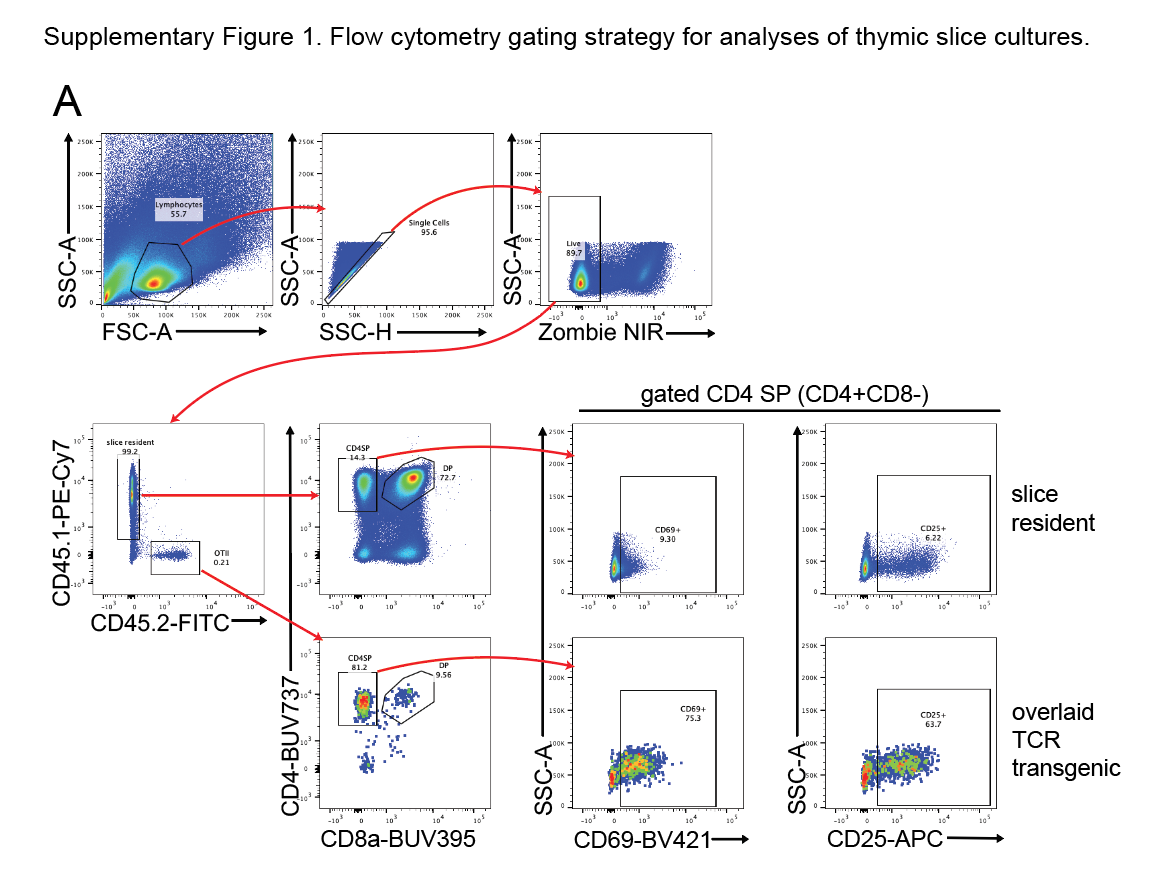

**Supplementary Figure 1. Flow cytometry gating strategy for analyses of thymic slice cultures.**

Representative flow plots displaying gating strategy for identifying TCR transgenic thymocytes in thymic slices. Samples were gated to include lymphocytes and exclude doublets and dead cells (Zombie NIR viability dye positive cells), and then gates were drawn to identify congenically distinct slice resident (CD45.1+) or overlaid TCR transgenic (CD45.2+) cells. Polyclonal slice resident (top) and overlaid TCR transgenic (bottom) CD4 SP (CD4+CD8-) thymocytes were then assessed for expression of TCR activation markers by gating on side scatter versus each marker, such as CD25 and CD69 shown here.

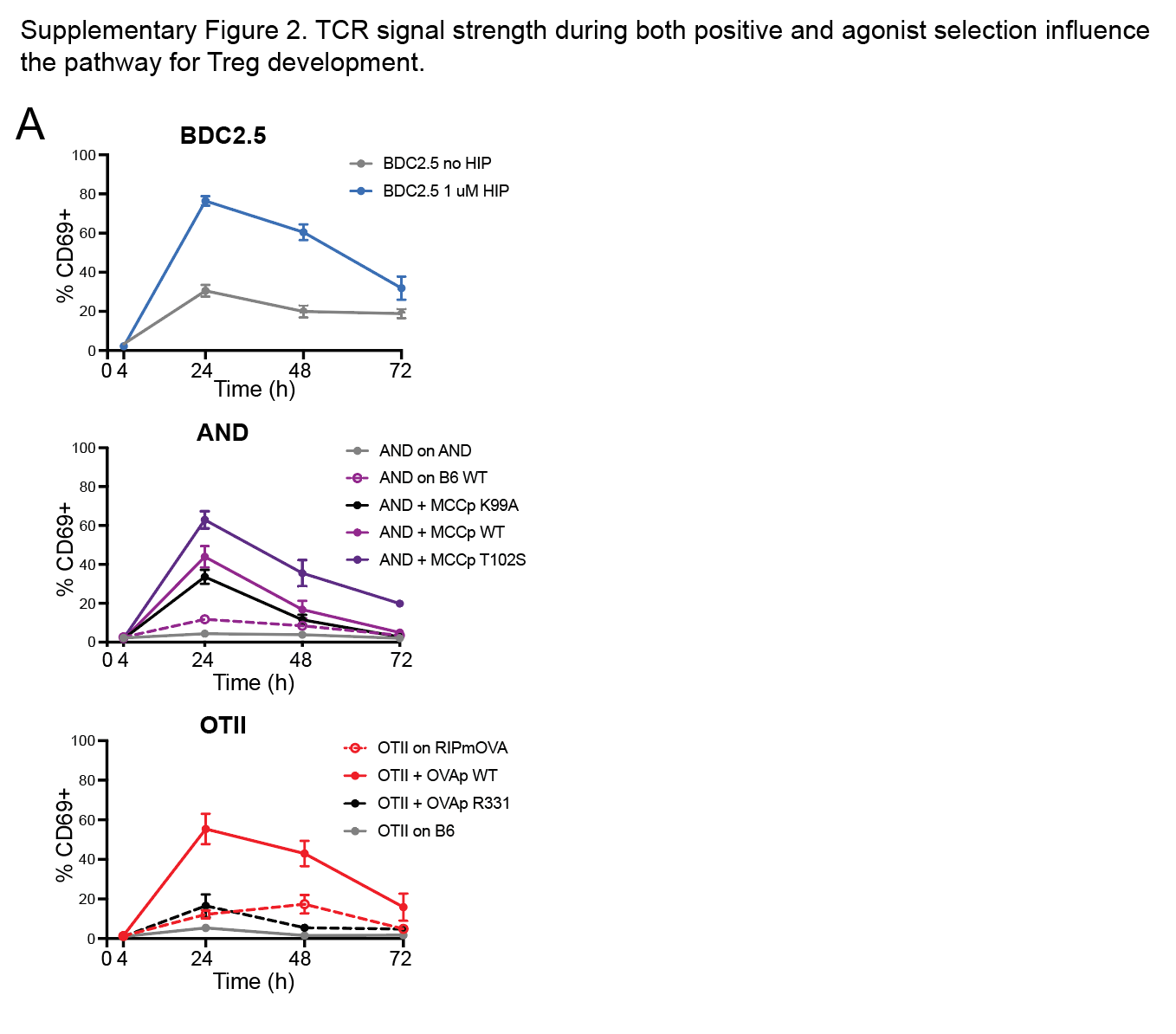

**Supplementary Figure 2. TCR signal strength during both positive and agonist selection influence the pathway for Treg development.**

A. Thymocytes from the indicated TCR transgenic were overlaid onto thymic slices using the stimulation conditions described in Fig. 2. Plots show the % CD69+ out of gated transgenic CD4 SP at the indicated times. Vertical error bars represent the SEM.

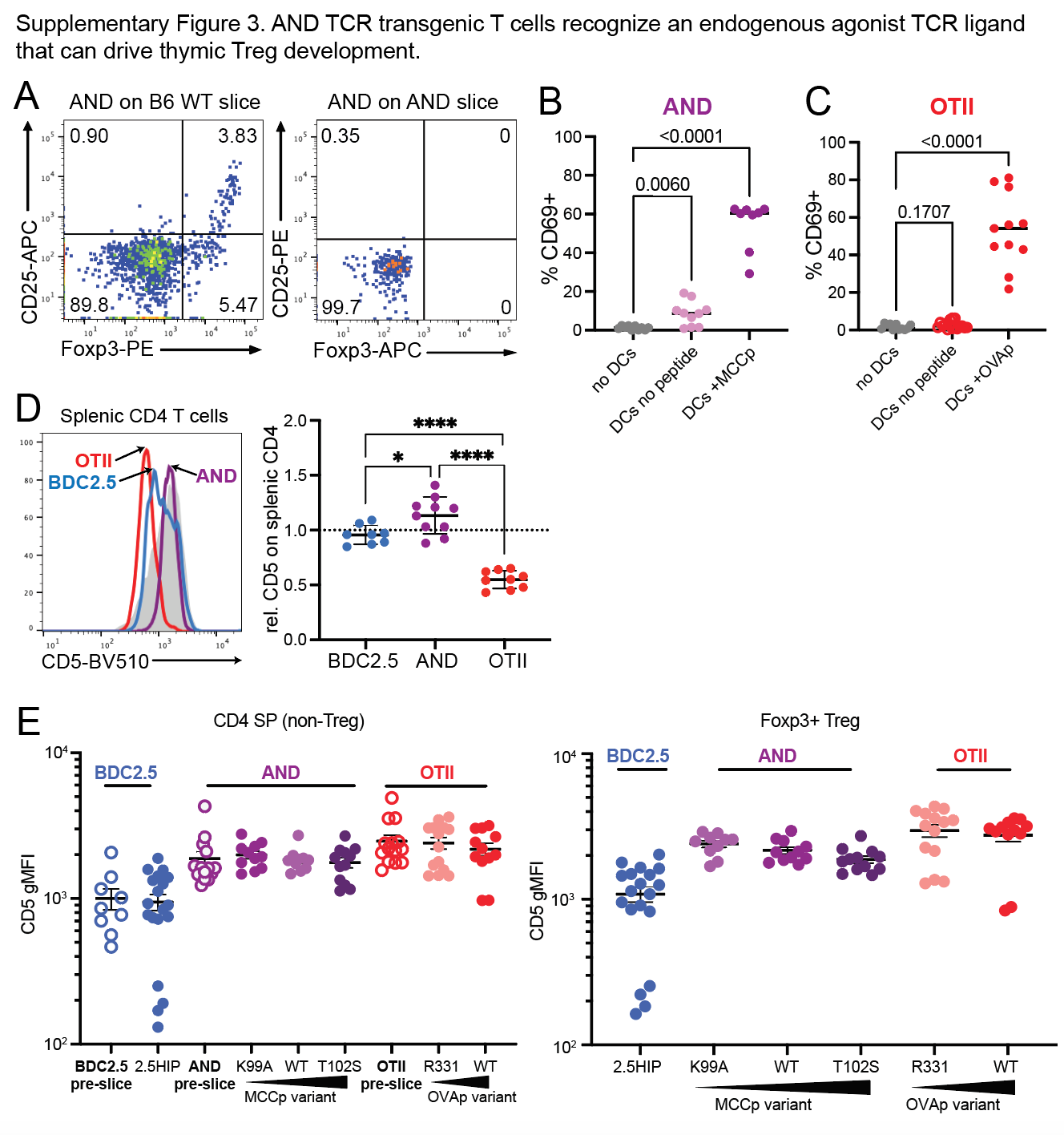

**Supplementary Figure 3. AND TCR transgenic T cells recognize an endogenous agonist TCR ligand.**

A. AND thymocytes give rise to Tregs on thymic slices from B6 WT, but not AND TCR transgenic mice. Flow plots showing CD25 and Foxp3 expression of AND CD4 SP thymocytes after 72 hours of culture on thymic slices from either B6 WT (left) or AND (right) mice. Data are representative of six experiments.

B,C. AND, but not OTII thymocytes respond to BMDC without added peptide. Thymocytes from AND (B) or OTII (C) TCR transgenic mice were cultured with or without BMDC and peptide for 24 hours and analyzed by flow cytometry. Plots show the % CD69+ out of gated CD4 SP. For AND thymocytes, F_1_ B6 WT x B10.BR BMDCs were used with or without 1 μM MCCp (B) and for OTII thymocytes, B6 WT BMDCs were used with or without 1 μM OVAp (C). For statistical analysis, one way analysis of variance (ANOVA) with multiple comparisons was used.

D. CD5 levels on splenic naive CD4 T cells (gated CD4+ CD44-) from the indicated TCR transgenic mice were determined by flow cytometry. Gray histograms are CD5 levels on CD4 T cells from B6 WT. CD5 gMFI for each sample was normalized to CD5 levels on equivalent WT cell population analyzed in the same experiment. Plots show representative flow cytometry plots (left) or compiled data (right). Six separate experiments with 1-2 mice each were performed, and each dot represents an individual mouse. Average values are indicated by horizontal bars, and range marks indicate SD. For statistical analysis, * = P <0.05, ****=P<0.0001 (one way analysis of variance (ANOVA) with multiple comparisons).

E. CD5 levels on BDC2.5, AND, or OTII thymocytes cultured on thymic slices for 72h using the stimulation conditions described in Fig. 2. Plots show the CD5 gMFI for each sample under each condition. The left graph shows CD5 levels on CD4 SP Foxp3- non-Tregs from each TCR transgenic model, and the right graph shows CD5 levels on CD4 SP Foxp3+ Tregs. On the left graph, open circles show the CD5 levels on BDC2.5, AND, or OTII thymocytes before culture on thymic slices for comparison (similar data shown in Fig 2A).

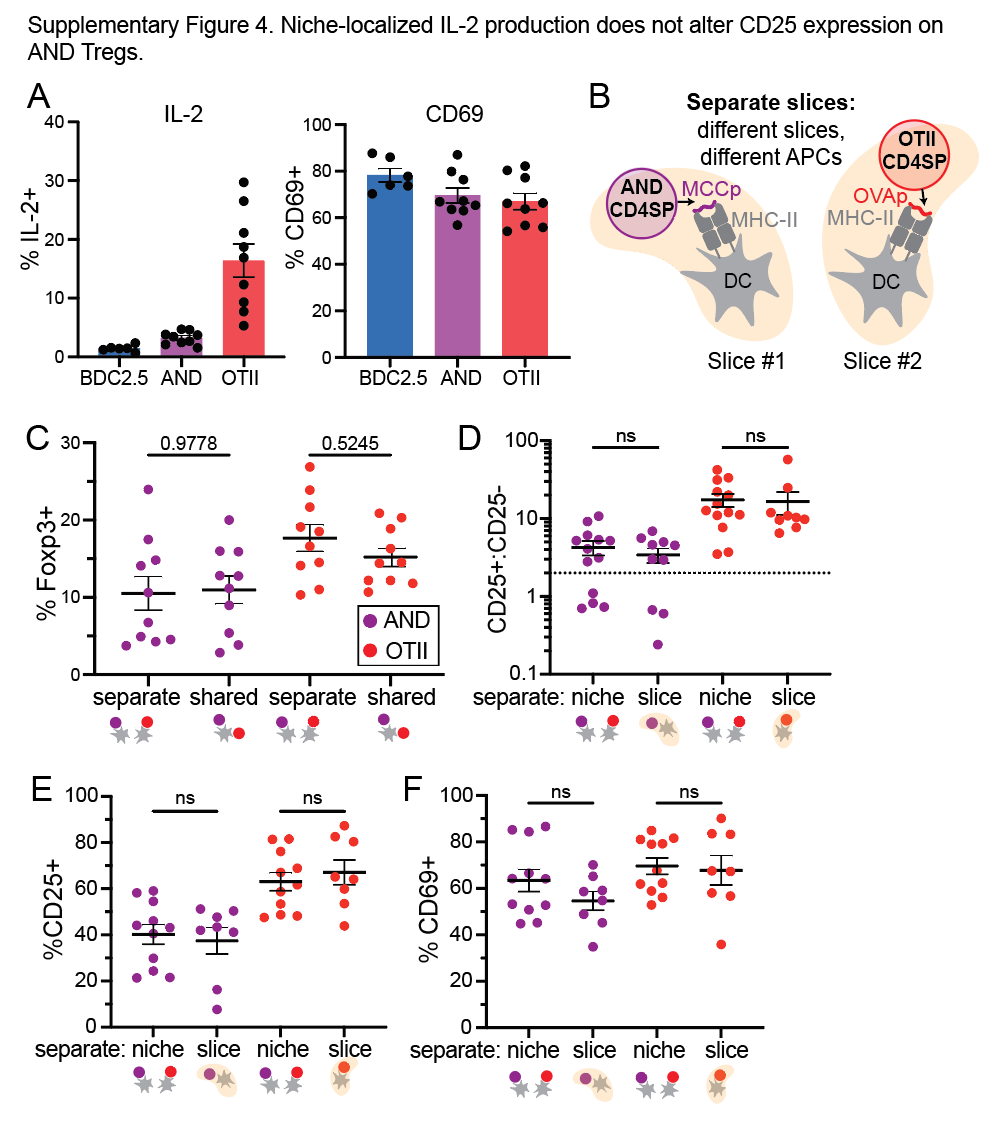

**Supplementary Figure 4. Niche-localized IL-2 production does not alter CD25 expression on AND Tregs.**

A. TCR transgenic thymocytes were stimulated *in vitro* with BMDC loaded with agonist peptide for 16 hours, and IL-2 and CD69 expression on TCR transgenic CD4 SP was determined by flow cytometry. Data is pooled from at least three experiments, and each symbol represents a replicate sample.

B. Schematic of separate slice controls for the shared niche experiment described in Fig. 5.

C. AND and OTII thymocytes were cultured on thymic slices along with peptide loaded BMDC from F_1_ (B6xB10.BR) mice. After 72 hours the total % Foxp3+ of CD4 SP was quantified to assess Treg development from AND or OTII donor populations. Shared and separate niche conditions were as described in Fig. 5, and data from Fig. 5 is included for comparison.

D,E, F. AND and OTII thymocytes were separately cultured on thymic slices along with peptide loaded BMDC from F_1_ (B6xB10.BR) mice (separate slice condition). Separate niche condition data from Fig. 5 D, E are shown for comparison. Samples were analyzed after 72 hours to assess CD25+ and CD25- Treg development (D) or 24 hours to assess CD25 and CD69 expression (E,F) from AND or OTII donor populations. Data is represented as the average +/- SEM. For statistical analysis, one-way ANOVA with multiple comparisons was performed.

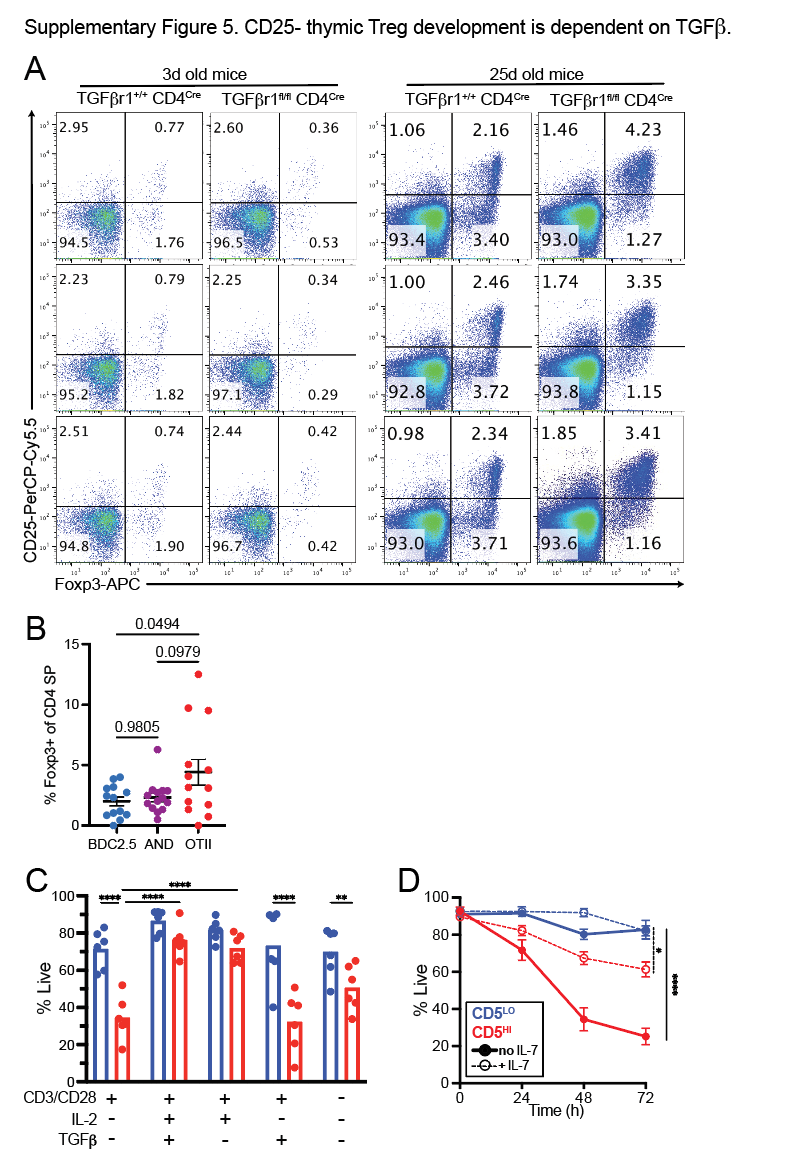

**Supplementary Figure 5. CD25- thymic Treg development is dependent on TGFB.**

A. Representative flow plots showing CD25 and Foxp3 expression of thymic Tregs from TGFβr1^fl/fl^ x CD4^Cre^ mice (control animals were TGFβr1^+/+^ x CD4^Cre^) were analyzed. Mice age 3d (left side) and 25d (right side) were used.

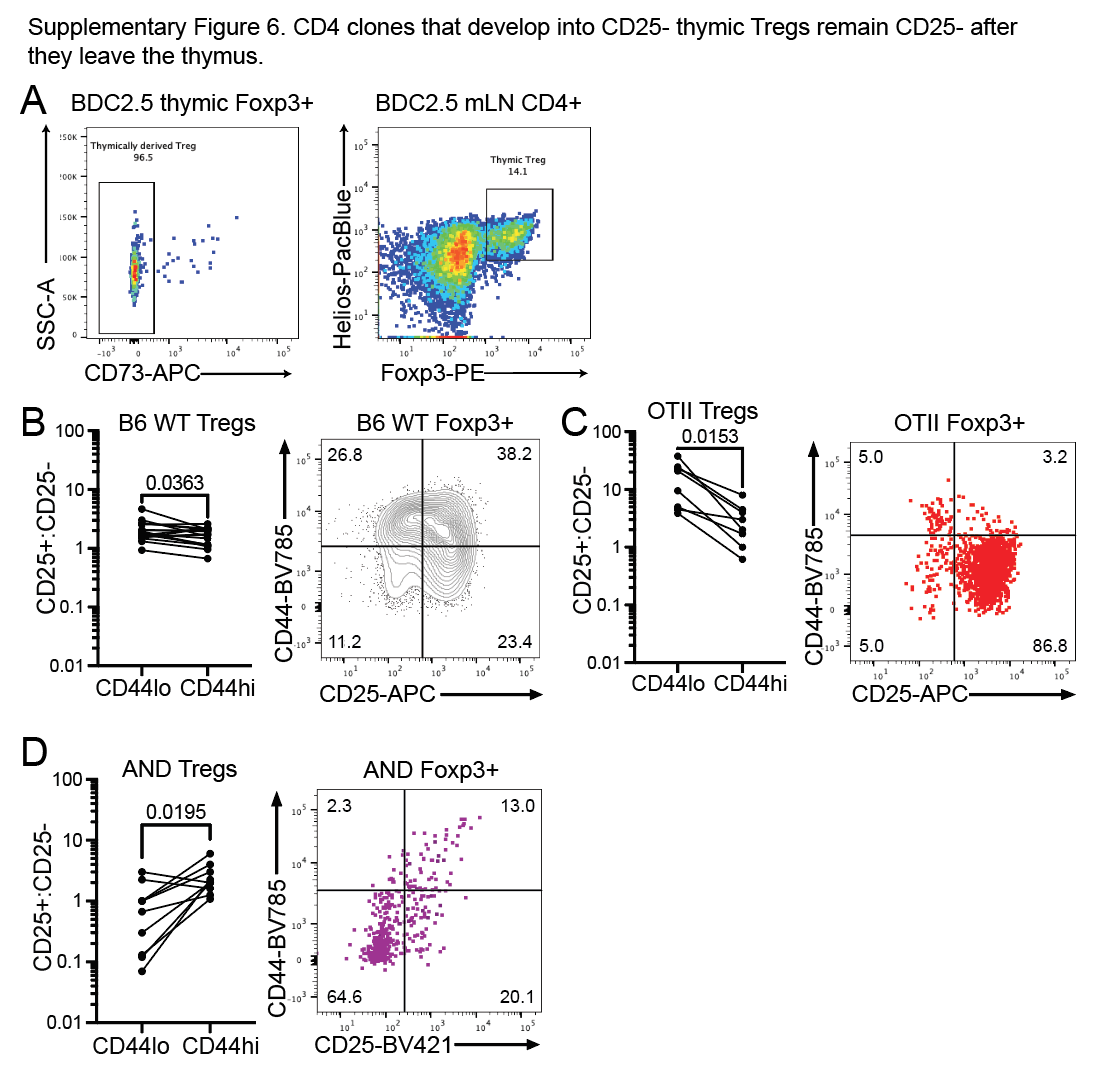

**Supplementary Figure 5. CD25- thymic Tregs remain CD25- as peripheral central Tregs.**

A. Thymus and lymph node data from neonatal BDC2.5 mice injected with Ab-2.5HIP. Right panel shows CD73 expression in Foxp3+ CD4 SP thymocytes. Left panel shows Helios and Foxp3 expression in CD4 T cells from mesenteric lymph nodes.

B-D. Spleen and lymph node data from the low frequency TCR transgenic mice described in Fig. 6A-C were analyzed to relate CD44 and CD25 expression on Tregs from B6 WT donors (B), OTII donors (C) or AND donors (D). Right panels show CD25 vs. CD44 expression on Tregs from a representative B6 WT sample (B) or multiple concatenated samples from TCR transgenic samples (C,D). Left panels show the ratio of CD25+/CD25- for CD44^LO^ or CD44^HI^ Tregs present in pooled lymph nodes and spleen of chimeric mice. Lines indicate pairwise comparisons of CD44^LO^ and CD44^HI^ Tregs from the same sample. For statistical analysis, paired t-test was used.

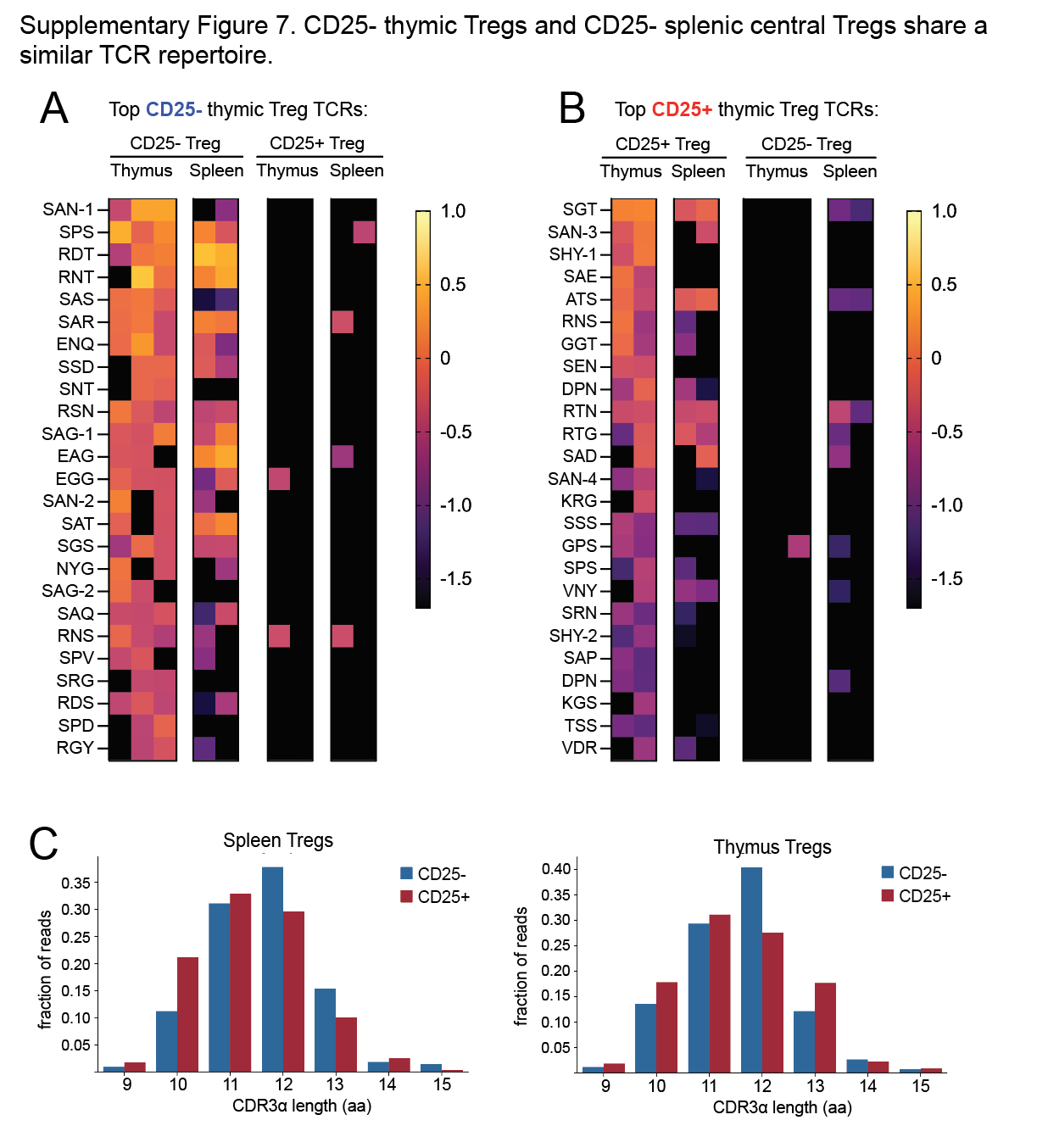

**Supplementary Figure 7. CD25- thymic Tregs and CD25- splenic central Tregs share a similar TCR repertoire.**

A, B. The frequency of the 25 most abundant recurring TCRs from CD25- thymic Tregs (A) or CD25+ thymic Tregs (B) from limited TCR diversity mice (described in Fig 6) plotted as a log_10_ value in heat map form. In A, the frequencies of the 25 most prevalent CD25- thymic Treg TCRs (leftmost plot) are shown, as well as the frequencies of those same TCRs in the CD25- splenic, CD25+ thymic, and CD25+ splenic Treg repertoires. B is similar to A except the frequencies of the 25 most abundant recurring CD25+ thymic Treg TCRs are shown.

C. CDR3⍺ length of splenic (top) or thymic (bottom) CD25+ or – Tregs represented as the fraction of reads corresponding to each CDR3⍺ length between 9-15 amino acids (aa).

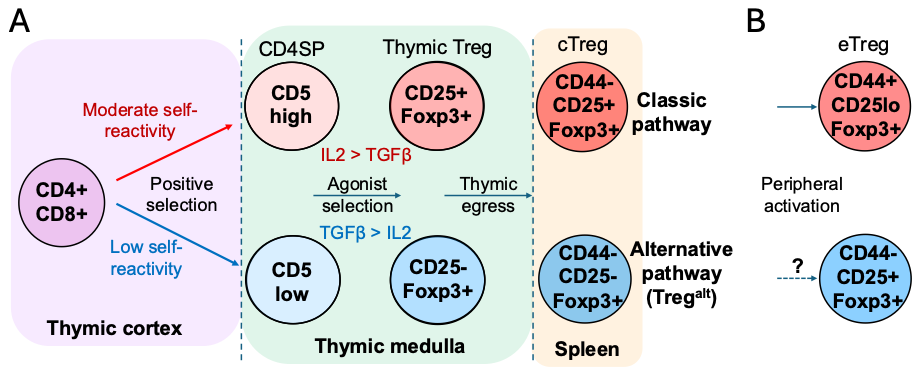

**Supplementary Fig. 8. A working model for thymic development of classic versus alternative (Treg^alt^) regulatory T cells.**

A. During positive selection thymocytes undergo functional tuning based on their level of reactivity to cortical self-peptide MHC, resulting in CD4 SP thymocytes with a range of responsiveness to cytokines and antigen, (Palmer 2011, Cho/Sprent, Persaud 2014, Mandl 2013 and 2026). This results in divergent responses to encounter with high affinity (agonist) medullary self-ligands and cytokines, leading to distinct Treg fates, depending on the strength of signal experienced during positive selection. Specifically, CD4 SP with moderate reactivity to cortical self (CD5^HI^) tend to upregulate CD25 and depend primarily on IL-2 to promote Treg development, whereas CD4 SP with low reactivity to cortical self (CD5^LO^) are predisposed to upregulate Foxp3 without CD25 induction, and depend primarily on TGFβ. After thymic egress, CD25-Tregs (Treg^alt^) retain their CD25- phenotype as central (CD44^LO^) Tregs (cTreg). B. After they leave the thymus cTregs can undergo further activation in the periphery to give rise to CD44^HI^ (effector) Tregs (eTreg). While classic Tregs tend to downregulate CD25 upon peripheral activation (Smigiel 2014), we speculate that Treg^alt^ cells can transiently induce CD25 upon activation, similarly to conventional T cells.
